# Binding of the integrated stress response inhibitor, ISRIB, reveals a regulatory site in the nucleotide exchange factor, elF2B

**DOI:** 10.1101/224824

**Authors:** Alisa F. Zyryanova, Félix Weis, Alexandre Faille, Akeel Abo Alard, Ana Crespillo-Casado, Heather P. Harding, Felicity Allen, Leopold Parts, Christophe Fromont, Peter M. Fischer, Alan J. Warren, David Ron

## Abstract

The Integrated Stress Response (ISR) is a conserved eukaryotic translational and transcriptional program implicated in mammalian metabolism, memory and immunity. The ISR is mediated by stress-induced phosphorylation of translation initiation factor 2 (eIF2) that attenuates the guanine nucleotide exchange factor eIF2B. A chemical inhibitor of the ISR, ISRIB, a bis-O-arylglycolamide, reverses the attenuation of eIF2B by phosphorylated eIF2, protecting mice from neurodegeneration and traumatic brain injury. We report on a cryo-electron microscopy-based structure of ISRIB-bound human eIF2B revealing an ISRIB-binding pocket at the interface between the β and δ regulatory subunits. CRISPR/Cas9 mutagenesis of residues lining this pocket altered the hierarchical cellular response to ISRIB congeners *in vivo* and ISRIB-binding *in vitro*, thus providing chemogenetic support for the functional relevance of ISRIB binding at a distance from known eIF2-eIF2B interaction sites. Our findings point to a hitherto unexpected allosteric site in the eIF2B decamer exploited by ISRIB to regulate translation.

## Introduction

Stress-induced phosphorylation of the α subunit of eukaryotic translation initiation factor 2 (eIF2α) is a highly conserved mechanism for regulating translation initiation (reviewed in Sonenberg and Hinnebusch, 2009). The resulting attenuation of global protein synthesis and the activation of translation from a rare subset of mRNAs that encode potent transcription factors is the basis of an Integrated Stress Response (ISR) (Dever et al., 1992; Harding et al., 2003). Whilst the ISR has important homeostatic functions that increase fitness, in some circumstances a benefit arises from attenuated signalling in the ISR (reviewed in Pakos-Zebrucka et al., 2016). This failure of homeostasis has encouraged the search for ISR inhibitors, which culminated in the discovery of ISRIB, the first small molecule ISR inhibitor (Sidrauski et al., 2013; Sidrauski et al., 2015a). ISRIB has proven efficacious in certain mouse models of neurodegeneration (Halliday et al., 2015), traumatic brain injury (Chou et al., 2017) and even as a memory-enhancing drug in normal rats (Sidrauski et al., 2013).

ISR inhibition in ISRIB-treated cells is observed despite elevated levels of eIF2(αP) (Sidrauski et al., 2013), indicating that ISRIB’s site of action lies downstream of the stress-induced kinases that phosphorylate eIF2α. The cellular target of eIF2(αP) is eIF2B, a large protein complex that possesses guanine nucleotide exchange (GEF) activity directed towards eIF2 (Panniers and Henshaw, 1983). The interaction of eIF2(αP) with eIF2B attenuates the recycling of eIF2 to its active, GTP-bound form, thereby attenuating rates of translation initiation on most mRNAs (Siekierka et al., 1981; Clemens et al., 1982) whilst enhancing initiation on rare mRNAs with peculiar 5’ untranslated regions (Mueller and Hinnebusch, 1986; Harding et al., 2000). Thus the ISR may be monitored by the activity of its target genes.

Genetic and structural observations indicate that the inhibitory effect of eIF2(αP) on eIF2B arises from the engagement by the α subunit of eIF2 at a cavity formed by the convergence of the α, β and δ regulatory subunits of the eIF2B decamer (Vazquez de Aldana and Hinnebusch, 1994; Pavitt et al., 1997; Kashiwagi et al., 2016; Bogorad et al., 2017). This regulatory site is distant from the catalytic γε subcomplex of eIF2B and its consecutive Asn-Phe residues that engage the nucleotide binding γ subunit of eIF2 to effect nucleotide exchange (Kashiwagi et al., 2016; Kashiwagi et al., 2017).

*In vitro*, addition of ISRIB accelerates eIF2B-mediated GDP dissociation from purified eIF2 (Sekine et al., 2015; Sidrauski et al., 2015b) and somatic cell genetic experiments have revealed that targeting eIF2B’s δ regulatory subunit can impart ISRIB resistance to cells (Sekine et al., 2015; Sidrauski et al., 2015b). An interaction of ISRIB with eIF2B is also consistent with the finding that the eIF2B decamer isolated from ISRIB-treated cells (or cell lysates) is more stable in a density gradient than eIF2B from untreated cells (Sidrauski et al., 2015b’, figure 3 therein). However, the ISRIB-resistant mutations identified to date cluster at a distance from both the regulatory site engaged by eIF2(αP) and the catalytic site engaged by eIF2γ (Kashiwagi et al., 2016; Kashiwagi et al., 2017). Thus, whilst the bulk of the evidence suggests that ISRIB binds eIF2B to regulate its activity, indirect modes of action are not excluded. Here we apply biophysical, structural and chemogenetic methods to reveal the presence of an ISRIB-binding pocket in eIF2B and provide insight into ISRIB’s mode of action.

## Results

### ISRIB binds directly to purified mammalian eIF2B

In keeping with previously-published observations (Sidrauski et al., 2015b), a stabilizing effect of ISRIB was observed on the eIF2B complex derived from HeLa cells in which the endogenous eIF2B β subunit was tagged (Figure 1A). This tagged complex was purified from intact cells (Figure 1B) and used to further study ISRIB’s interaction with eIF2B.

**Figure 1.**
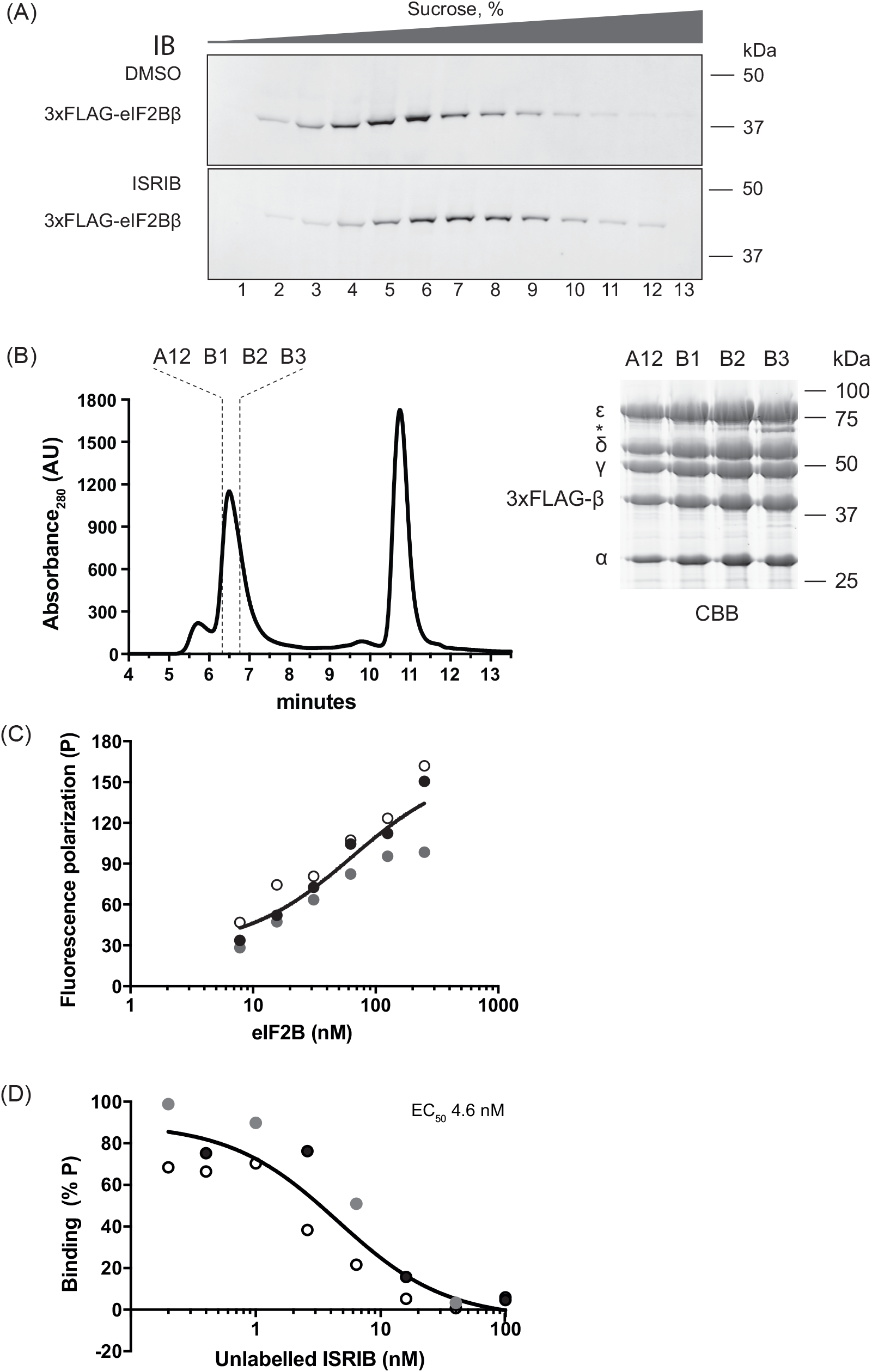
ISRIB binds to purified eIF2B. (A) Immunoblot of 3xFLAG-tagged eIF2Bβ detected with anti-FLAG M2 antibodies in HeLa cell lysates that were either treated with DMSO (top panel) or ISRIB (200 nM) (bottom panel) and resolved on a 5-20% sucrose density gradient. The observed shift of 3xFLAG-eIF2Bβ signal towards HMW fractions reflects stabilizing effect of ISRIB on the protein complex (compare top and bottom panels) (performed once, confirming the findings of Sidrauski et al., 2015b) (B) Chromatogram of endogenous human eIF2B purified from HeLa cells with 3xFLAG-tag (introduced by homologous recombination into the *EIF2B2* locus), resolved by size exclusion chromatography on a SEC-3 300 Å chromatography column (left panel). The indicated fractions were analyzed on a Coomassie-stained SDS-PAGE gel (right panel, the asterisks * indicates PRMT5 – a contaminant) and used in structural analysis. (C) A plot of the fluorescence polarization (FP) signal arising from FAM-labelled ISRIB congener (AAA2-101) (2.5 nM) as a function of the concentration of eIF2B in the sample. Concentrations of eIF2B are represented on a log_10_ scale. Curve fitting was generated using “agonist vs. response” function on GraphPad Prism, shown are values of three independently-acquired measurements. (D) A plot of the FP signal arising from samples with FAM-labelled AAA2-101 (2.5 nM) bound to purified human eIF2B (30 nM) in the presence of the indicated concentration of unlabelled trans-ISRIB introduced as a competitor. Concentrations of ISRIB are represented on a log_10_ scale. Curve fitting and EC_50_ was generated using “agonist vs. response” function on GraphPad Prism, shown are values of three independently-acquired measurements.

To observe ISRIB’s interaction with eIF2B directly, we designed a fluorescently labelled derivative (AAA2-101) based on known structure–activity relationships of ISRIB derivatives (Hearn et al., 2016) (Figure 1-supplement 1A). These show that major modifications of the central *trans-cyclohexyl* group and the glycolamide linkages are poorly tolerated, whereas changes to the terminal aryl groups in some cases maintain or enhance activity. We observed that tethering of the glycolamide C^α^ of one of the cyclohexyl substituents to the neighbouring phenyl in the form of a chromane group (AAA1-084, Figure 1-supplement 1A) retained activity and was only 8.5-fold less active than trans-ISRIB in cells (Figure 1-supplement 1B). Replacement of the chromane with the dihydrobenzoxazine system afforded a compound (AAA1-075B, Figure 1-supplement 1A) only 3-fold less active than trans-ISRIB in cells (Figure 1-supplement 1B). The aniline function in this compound was then used to elaborate the fluorescently labelled compound AAA2-101, which contains a flexible linker terminating in a 6-carboxyfluorescein moiety (Figure 1-supplement 1A).

Using this compound we measured the effect of added purified eIF2B on the fluorescence polarization (FP) signal. FP increased with increasing concentrations of eIF2B (Figure 1C). At the concentrations of eIF2B available for testing, the FP signal was not saturated, thus we have been unable to extract a reliable dissociation constant from this assay. However, unlabelled ISRIB effectively competed for eIF2B in the FP assay, with an EC_50_ in the nanomolar range, as observed for ISRIB action in cells (Figure 1-supplement 1B), whilst less active congeners competed less successfully (Figure 1D and Figure 1-supplement 1C). The FP assay thus likely reports on engagement of the FAM-labelled ISRIB derivative at a site(s) on eIF2B that is relevant to ISRIB action.

### Cryo-EM reveals an ISRIB binding pocket in eIF2B

The structures of several eIF2B-related complexes have been solved by X-ray crystallography: the human α_2_ dimer (Hiyama et al., 2009), the *C. thermophilum* (βδ)_2_ regulatory sub-complexes (Kuhle et al., 2015) and the complete *S. pombe* eIF2B decamer (Kashiwagi et al., 2016). However, as fission yeast eIF2B does not respond to ISRIB and as there is an inherent uncertainty if the partial complexes would bind ISRIB, we elected to solve the structure of the ISRIB-bound human eIF2B complex by single particle cryo-electron microscopy (cryo-EM).

We purified endogenous human eIF2B from HeLa cell lysates in the presence of ISRIB and determined the structure of the ISRIB-eIF2B complex at an overall resolution of 4.1-Å (Figure 2A, Figure 2- supplement 1 and table 1). Within the β and δ regulatory core, in the most central part of the map, protein side chains are clearly resolved, resulting in a near complete atomic model (Figure 2B, C and Figure 2- supplement 2A & 2B). The local resolution of the map varies from 3.7 Å in the core to >6 Å at the periphery (Figure 2-supplement 2A). In particular, the resolution of the γ and ε human catalytic subcomplex was lower compared with the regulatory core, consistent with flexibility of this region in both human and yeast eIF2B (Kashiwagi et al., 2016). Similar to the *S. pombe* eIF2B crystal structure, we were unable to resolve the catalytically important C-terminal HEAT domain of the ε subunit in our cryo-EM map.

Due to lack of sufficient high-quality cryo-EM images of an apo-eIF2B complex, we were unable to calculate a difference map of eIF2B with and without ISRIB. However, a nearly continuous density with a shape and size of a single ISRIB molecule was conspicuously present at the interface of the β and δ regulatory subunits. The quality of the map in this position provided sufficient detail to confidently model the ISRIB molecule (Figure 2C). The ISRIB binding pocket is located at the plane of symmetry between the β and δ subunits. In the central part of the pocket, residue H188 of the β subunit is notable, as its side chain is positioned in the vicinity of the essential carbonyl moiety of ISRIB (Hearn et al., 2016). Residue N162 of the β subunit also stabilises the diaminecyclohexane moiety of ISRIB, possibly through hydrogen bonding interactions. More distally in the pocket, various residue side chains – namely δL179 (the human counterpart of hamster δL180), δF452, δL485, δL487, βV164, βI190, βT215, and βM217 – form the hydrophobic end of the symmetrical pocket that accommodates the aryl groups of ISRIB (Figure 2-supplement 2C).

**Figure 2.**
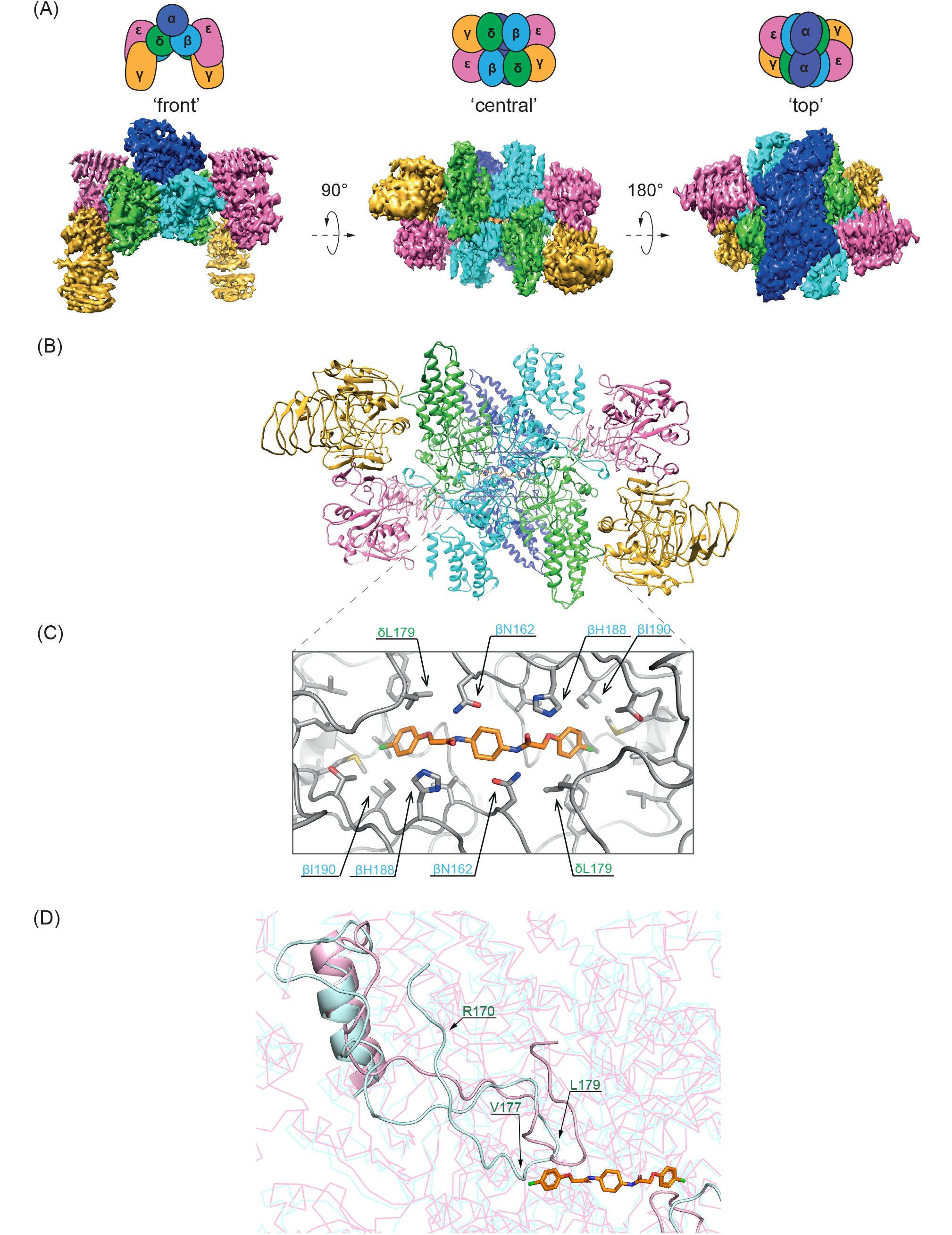
Structure of the ISRIB-bound human eIF2B. (A) Representative views of the unfiltered cryo-EM map of the ISRIB-bound decameric human eIF2B complex. Density is colored according to the subunit architecture indicated in the cartoons: α – blue, β – cyan, δ – green, γ – gold, ε – pink. (B) Ribbon representation of ISRIB-bound human eIF2B. ‘Central’ view of the (βδ)_2_ dimer interface reveals a density corresponding to a single molecule of ISRIB. (C) Close-up of the ‘central’ view showing the ISRIB binding site. An ISRIB molecule is docked in the density observed in the cavity at the (βδ)_2_ dimer interface. Residues contacting ISRIB in the central part of the pocket from the β (blue) and δ (green) subunits are indicated. ISRIB is represented in orange sticks. (D) Cartoon showing the divergence in orientation of the N-terminal portion of the ISRIB-bound human δ subunit (residues 167-204, in cyan) and yeast δ subunit (residues 114-142, from PDB: 5B04, in light magenta). Three residues: human δ R170, V177, L179 (important for ISRIB activity in cells Sekine et al., 2015) are indicated. ISRIB is in orange. Note the convergence of the structure at δ Proline 205 (human)/ δ Proline 143 (yeast), which cap the closely-aligned helix I of the human and yeast eIF2Bδ

The observation that a hamster *Eif2b4^L180F^* mutation (δL179 in the human) disrupts ISRIB action in cells (Sekine et al., 2015) is consistent with a clash between the bulkier side chain of phenylalanine and the bound ISRIB molecule. The ISRIB binding site identified in the map is also consistent with the lack of a corresponding density in the *S. pombe* apo-eIF2B map.

Overall the human and yeast structures are highly similar (r.m.s.d of 2.57 Å over 3049 alpha carbons) with no evidence of major domain movements that might be attributed to ISRIB binding in the human complex (Figure 2 –supplement 2D). The region corresponding to the N-terminal ~166 residues of the human δ subunit remains unresolved (in both yeast and human structures).

However, residues 167-204 of the human protein, which can be reliably placed in the density, assumed a very different conformation from the corresponding segment of the yeast δ subunit (Figure 2D). It is unclear if this difference reflects species-divergence in the structure of eIF2B, a crystallization-induced difference or is a consequence of ISRIB binding. It is noteworthy, however, that this divergent region emerges from the putative ISRIB density at the (βδ)_2_ interface and that it contains two residues, human eIF2B δ R170 and V177, whose mutation interferes with ISRIB action in hamster cells, δ R171 and V178 (reported in Sekine et al., 2015).

### Chemogenetic analysis of the putative ISRIB-binding pocket

To validate the mode of ISRIB binding revealed by the structural data (Figure 2C) we randomized the codons, encoding residues predicted by the model to line the ISRIB-binding pocket, in hope of eliciting a correlation between amino acid substitutions and ISRIB’s activity in such mutagenized cells. To achieve this, we used CRISPR/Cas9 to target the aforementioned *Eif2b2* locus of CHO-K1 cells and provided a repair template randomized at either *Eif2b2^N162^, Eif2b2^H188^* or *Eif2b2^I190^* codons (Figure 3A). Exposure to histidinol, an agent that activates the eIF2α kinase GCN2, induces an ISR, which can be tracked by a CHOP::GFP reporter in living cells (Sekine et al., 2015). In a first round of sorting, histidinol-treated, mutagenized pools of cells, were segregated by phenotypic selection into ISRIB sensitive [ISRIB^SEN^, CHOP::GFP inhibited (“GFP-dull”)], and ISRIB resistant [ISRIB^RES^, CHOP::GFP activated (“GFP-bright”)] classes. In a second round of sorting untreated populations from the ISRIB^SEN^ and ISRIB^RES^ pools were purged of GFP-bright cells that had acquired a constitutively active ISR phenotype (Figure 3B).

**Figure 3.**
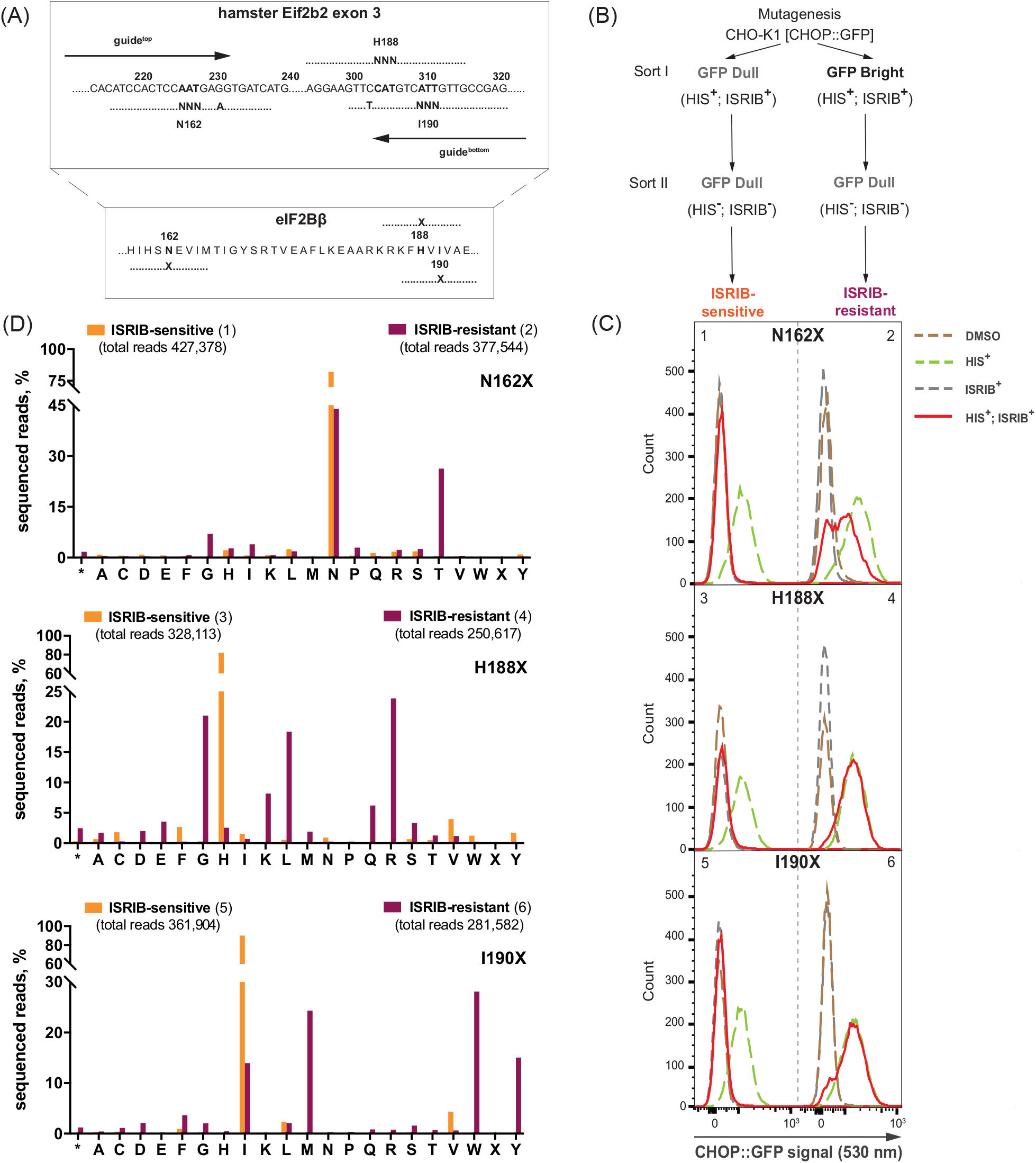
Structure-directed chemogenetic analysis of ISRIB binding to eIF2B. (A) Schema of the CRISPR/Cas9 directed mutagenesis of the hamster *Eif2b2* gene at the locus encoding ISRIB contact sites on the eIF2B β subunit revealed by the structure: N162, H188 or I190. Hamster *Eif2b2* exon 3 with and the two RNA guides directing Cas9 to mediate double strand break, and the aligned single-strand DNA repair templates. The randomized substitution of the original codon is indicated (NNN), as are the silent PAM-disrupting mutations. Nucleotide numbering corresponds to GeneBank accession NW_003613913.1. (B) Schema for selecting ISRIB-sensitive (ISRIB^SEN^) and ISRIB-resistant (ISRIB^RES^) cells (Sort I) and eliminating cells with constitutive ISR (Sort II) following CRISPR/Cas9 mutagenesis, based on the ISR-activated CHOP::GFP fluorescent reporter upon histidinol (HIS) treatment of CHO-K1 cells. (C) Histograms of CHOP::GFP activity in ISRIB^SEN^ (left panels) and ISRIB^RES^ (right panels) pools selected as indicated in the panel 2B (above). Cells were treated with HIS (0.5 mM), ISRIB (200 nM) or both. (D) A bar graph showing the distribution of residues identified at the indicated positions of mutagenized *Eif2b2*, analysed by the next generation sequencing (NGS). Shown is a number of sequenced reads for ISRIB^SEN^ (orange bars) and ISRIB^RES^ (plum bars) for each amino acid (* - stop codon, X – ambiguous sequence).

Using fluorescence activated cell sorting (FACS) we successfully isolated ISRIB^SEN^ cells, in which ISR induction by histidinol was readily counteracted by ISRIB, as well as ISRIB^RES^ cells, in which ISR induction was maintained upon ISRIB treatment (Figure 3C – compare left and right panels, red traces). To determine if the phenotypically-distinguished pools were enriched in different mutations, we subjected genomic DNA derived from each population to deep sequencing analysis (Figure 3D and table S2).

The ISRIB^RES^ pool of cells targeted at *Eif2b2^H188^* diverged dramatically from the parental sequence (Figure 3D – middle panel, plum bars). Of a total of 250,617 reads histidine was present in only 6,443 (2.6%), with arginine, glycine, leucine, lysine and glutamine, dominating (24%, 21%, 18%, 8.2% and 6.2%, respectively). Histidine was preserved in the ISRIB^SEN^ pool (269,253 of 328,113 reads, 82%), which was constituted largely of parental alleles that escaped targeting altogether and targeted alleles that had acquired a synonymous mutation (Figure 3D – middle panel, orange bars). These observations point to the importance of histidine at position 188 of the β subunit in mediating ISRIB sensitivity. The bias in a favour of certain substitutions in the ISRIB^RES^ pool likely arises by their ability to preserve eIF2B function whilst eliminating responsiveness to ISRIB.

The ISRIB^RES^ pool of cells targeted at *Eif2b2^I190^* was dominated by tryptophan, methionine and tyrosine (28%, 24% and 15%, respectively) consistent with a role for these bulky side chains in occluding the ISRIB binding pocket (Figure 3D – bottom panel, plum bars). This conclusion is supported by the observation that replacement by the smaller valine was relatively enriched in the ISRIB^SEN^ population (Figure 3D – bottom panel, orange bars).

Mutagenesis of *Eif2b2^N162^* was less successful in generating a pool of strongly ISRIB-resistant cells (Figure 3C, panel 2, red trace). This is not a reflection of failure of the homologous recombination strategy, as the incorporation of a silent PAM disrupting mutation enabled us to restrict the sequencing analysis to alleles that had successfully undergone homologous recombination. The high frequency with which the parental asparagine had been retained in both ISRIB^SEN^ and ISRIB^RES^ pools (82% and 44%, respectively) suggests that replacement of this residue is selected against (*Eif2b2* is an essential gene), with the one exception being acquisition of a threonine, which was greatly enriched (26%) in the ISRIB^RES^ pool (Figure 3D – top panel, plum bars).

While these features of the ISRIB^RES^ mutations suggest that they exert their effect by altering the character of an ISRIB-binding pocket, it is impossible to exclude an alternative possibility that they alter eIF2B structure so as to disrupt the communication between an ISRIB binding site that might be located elsewhere and the relevant catalytic activity of eIF2B. To examine this possibility, we tested pools of ISRIB^RES^ cells for residual responsiveness to ISRIB congeners, reasoning that a mechanism involving an altered binding site might alter the hierarchy of potency of ISRIB congeners, whereas a mechanism involving disruption of an allosteric signal is unlikely to do so.

Compounds AAA1-075B and AAA1-084 are 3- to 11-fold less potent than ISRIB in inhibiting the ISR of wildtype cells (Figure 1-supplement 1B, Figure 4A, figure 4 – supplement 1A). However, both compounds were relatively more potent than ISRIB in reversing the ISR of the ISRIB^RES^ pool of cells targeted at *Eif2b2^H188^*, attainting nearly complete inhibition when applied at micromolar concentration (Figure 4B). The faint, biphasic response of the mutant ISRIB^RES^ pool to ISRIB is reproducible, but currently lacking an explanation (Figure 4–supplement 1B). These observations point to a shift in the binding properties of the ISRIB pocket, induced by the mutations at position 188 of the β subunit.

**Figure 4.**
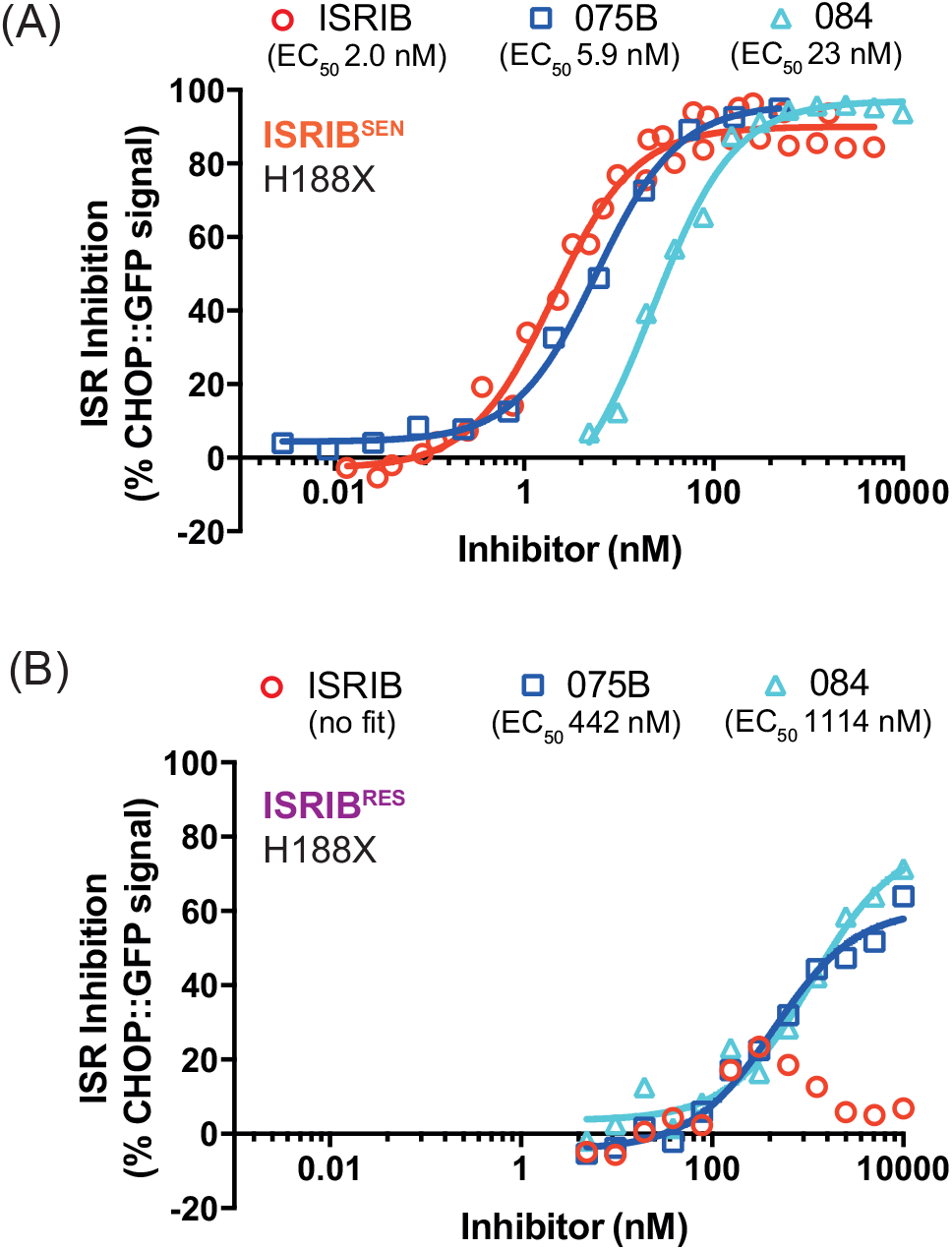
Mutations in Eif2b2 codon 188 (β^H188X^) alter selectivity for ISRIB congeners. Graphs showing inhibition of the ISR-activated CHOP::GFP signal (induced upon treatment with HIS, 0.5 mM) by ISRIB and two related derivatives, AAA1-075B (75B) and AAA1-084 (84), in ISRIB^SEN^ (A) and ISRIB^RES^ (B) mutant pools of *EIf2b2^H188X^*, revealed by flow cytometry. Shown is a representative from three independent experiments for each of the compounds (replicated in Fig. 4 – Sup. 1 A, B). Concentration of inhibitor is represented on a log_10_ scale. Curve fitting and EC_50_ was generated using “agonist vs. response” function on GraphPad Prism.

To consolidate the aforementioned findings, we exploited the diversity of ISRIB^RES^ mutations in the *Eif2b2^H188X^* population to select, by a new round of FACS (Figure 5A), for sub-pools that acquired sensitivity to AAA1-075B or AAA1-084, or retained their sensitivity to ISRIB (Figure 5B, compare continuous traces) and sequenced their *Eif2b2* alleles (Figure 5C and table S3). As expected, sorting for ISRIB sensitivity enriched, by over 20-fold for those rare wildtype H188 alleles that persisted the pool of ISRIB^RES^ *Eif2b2^H188X^* cells (Figure 5C – compare plum to orange bars). H188 was also somewhat enriched (about 5-fold) in the pools sorted for their sensitivity to AAA1-075B (075B^SEN^) or AAA1-084 (084^SEN^), but unlike the ISRIB^SEN^ these pools were also enriched for residues other than histidine (Figure 5C, blue and cyan bars). Importantly, selecting for sensitivity to these ISRIB congeners enriched for different residues than those found in the original ISRIB^RES^ pool: arginine, glycine and leucine were depleted and replaced by lysine, serine, alanine and threonine (Figure 5C, compare plum bars to blue and cyan bars).

**Figure 5.**
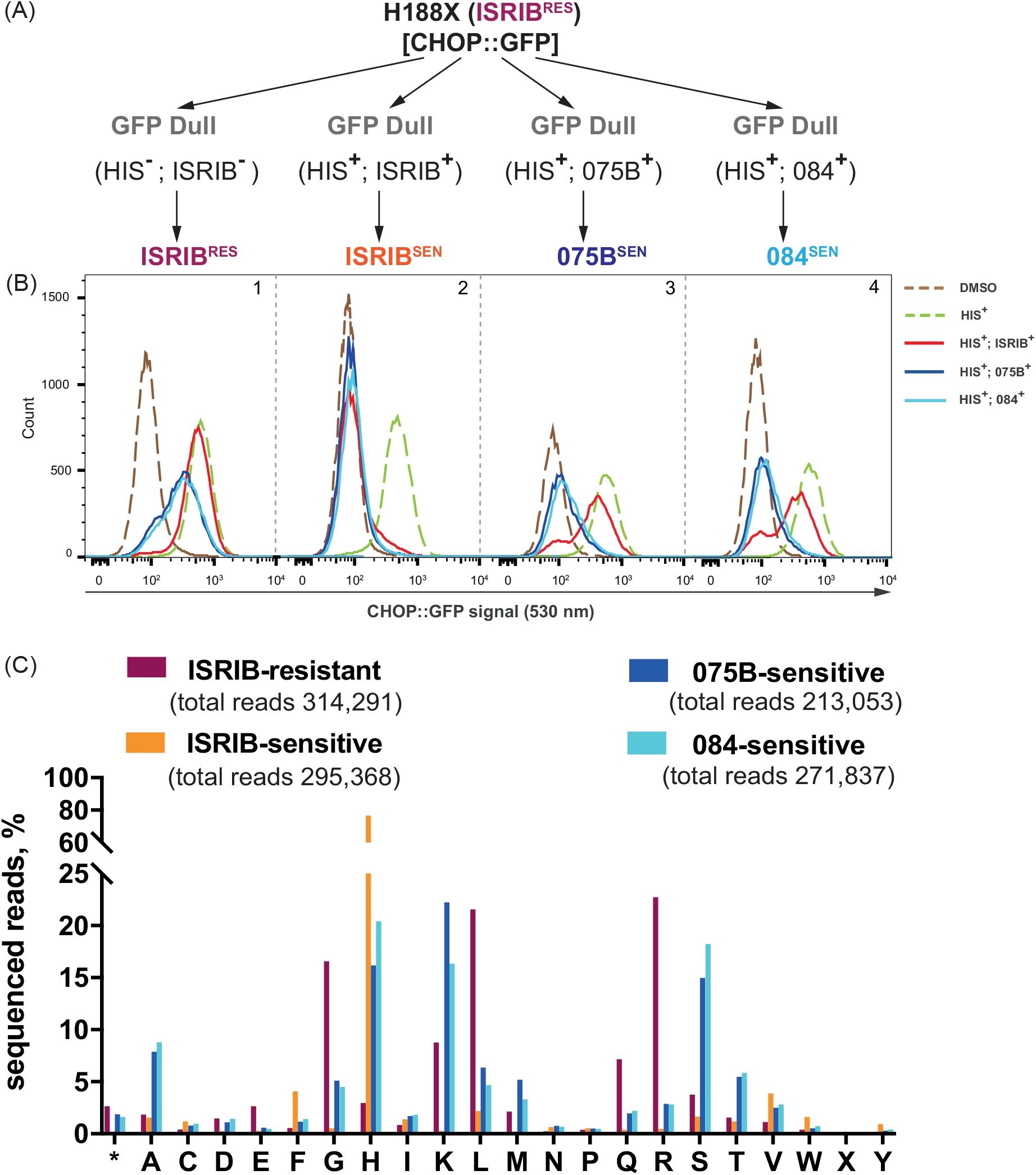
Sensitivity to ISRIB congeners selects for divergent palate of mutations in codon 188 of EIf2b2. (A) Schema for selecting ISRIB^RES^, ISRIB^SEN^, AAA1-075B sensitive (75B^SEN^) and AAA1-084 sensitive (84^SEN^) cells, based on activation of CHOP::GFP fluorescent reporter upon ISR induction in *Eif2b2^H188X^* mutant population. (B) Histograms of CHOP::GFP activity in ISRIB^RES^, ISRIB^SEN^, 75B^SEN^ and 84^SEN^ sub-pools selected from a population of originally ISRIB^RES^ cells selected as indicated in the panel 3A (above). Cells were treated with HIS (0.5 mM) ± ISRIB or its congeners (2.5 μM). (C) A bar graph showing the distribution of residues identified at codon 188 of *Eif2b2* analysed by NGS. Shown is a number of sequenced reads for ISRIB^RES^ (plum), ISRIB^SEN^ (orange), 75B^SEN^ (blue) and 84^SEN^ (cyan) for each amino acid (* - stop codon, X – ambiguous).

To examine the sensitivity to ISRIB congeners in individual clones of *Eif2b2^H188X^* mutant cells, we selected for study clones with unambiguous genotypes (Figure 6–supplement 1A). CHO cells that had undergone the aforementioned selection scheme but remained homozygous for the parental *Eif2b2^H188^* allele retained their responsiveness to ISRIB and AAA1-084. The responsiveness of the mutant clones to ISRIB was greatly enfeebled, noticeable in both a shift to the right in the concentration-response curves and in the magnitude of inhibition observed at the highest concentration of compound (Figure 6A, compare red traces across clones). The biphasic response to ISRIB, observed in the pools of mutagenized cells (Figure 4B), was also evident in the clonal populations. A similar shift to the right was noted in the response of the mutant clones to AAA1-084 but the inhibitory effect observed at the higher concentration was only slightly diminished (Figure 6A, compare blue traces).

**Figure 6.**
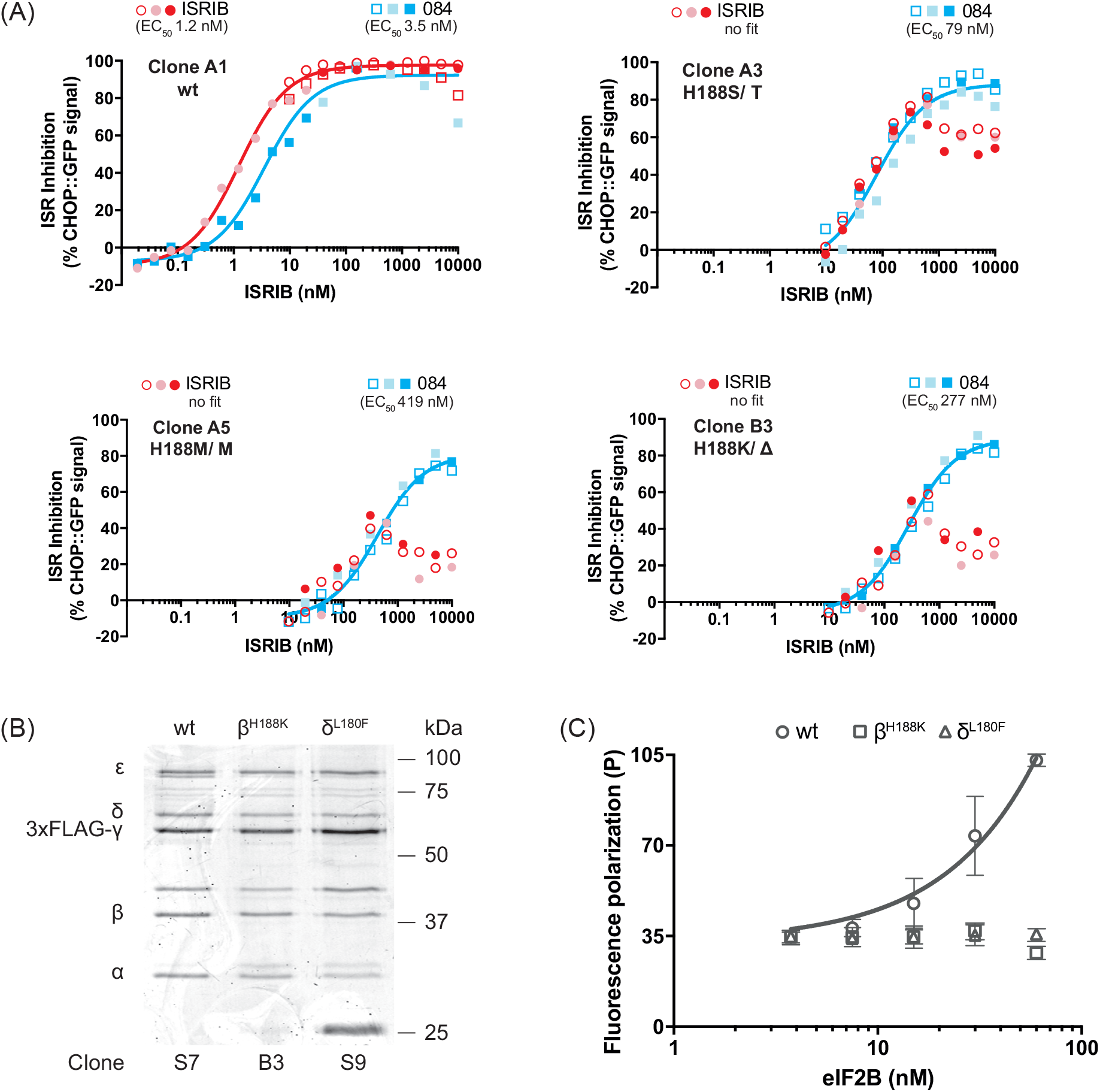
Mutations that impair ISRIB action in cells, impair ISRIB binding to eIF2B in vitro. (A) Graphs showing inhibition of the ISR-activated CHOP::GFP signal, induced upon treatment with HIS (0.5 mM), by ISRIB and AAA1-084 (084) in individual clones: A1 (parental, wt), A3 (H188S/T), A5 (H188M), B3 (H188K/Δ), isolated from 84^SEN^ pool (from Figure 5B). Shown is data from three independent experiments for each of the compounds. Concentration of inhibitor is represented on a log_10_ scale. Curves fitting and EC_50_ was generated using “agonist vs. response” function on GraphPad Prism. (B) Coomassie stained SDS-PAGE gel of endogenous hamster eIF2B purified from wildtype (wt, clone A1), *Eif2b2^H188K^* (clone B3) and *Eif2b4^L180F^* (clone S9, from Sekine et al., 2015) CHO cells via a 3xFLAG-tag knocked into *Eif2b3* locus (encoding the γ subunit). (C) A plot of the FP signal arising from FAM-labelled AAA2-101 (2.5 nM) as a function of the concentration of wildtype (wt) or mutant eIF2B (δ^L180F^ or β^H188K^) in the sample. Shown are mean ± SD (n=3). Concentrations of eIF2B are represented on a log_10_ scale. The fitting curve and EC_50_ was generated using “agonist vs. response” function on GraphPad Prism.

To address the effect of ISRIB-resistant mutations in eIF2B on binding of the FAM-labelled AAA2-101, we purified eIF2B from wildtype, *Eif2b4^L180F^* and *Eif2b2^H188K^* CHO cells by exploiting a 3×FLAG-tag knocked into the endogenous eIF2B γ subunit (Figure 6B). The wildtype eIF2B gave rise to a conspicuous concentration-dependent FP signal in the presence of a FAM-labelled AAA2-101 (Figure 6C, circular pictograms). Validity of this FP signal was confirmed by competition with unlabelled ISRIB (Figure 6 –supplement 1B). However eIF2B purified from the mutant cells failed to give rise to an FP signal (Figure 6C, square and triangle pictograms), thereby establishing a correlation between ISRIB resistance in cells and defective ISRIB binding *in vitro* (Figure 6A, C and Figure 6 – supplement 1C).

## Discussion

We have identified the ISRIB binding site at the core of the regulatory complex by determining the cryo-EM structure of a human eIF2B-ISRIB complex. The residues that contact ISRIB can be readily identified in the experimentally-derived density map. Mutation of these residues leads to loss of sensitivity to ISRIB in cultured cells and to loss of ISRIB binding *in vitro*, but the same mutations exert less of an enfeebling effect on the cellular response to certain ISRIB congeners. Together these structural, biophysical and chemogenetic findings point to a role for ligand engagement at the aforementioned pocket in ISRIB action.

The ISRIB binding pocket straddles the two-fold axis of symmetry of the core regulatory complex and a single molecule of ISRIB appears to engage the same residues from opposing protomers of the (βδ)_2_ dimer of dimers. This feature fits well with ISRIB’s own symmetry and also nicely explains the ability of ISRIB to stabilize the eIF2B decamer *in vitro*.

The overall structure of ISRIB-ligated human eIF2B is similar to that of *S. pombe* eIF2B, crystallized in the absence of ISRIB. This finding argues against large domain movements as the basis of ISRIB action. However, as we do not have information on the *apo* structure of human eIF2B, we cannot exclude the possibility that ISRIB stabilizes an active conformation that was also fortuitously assumed by yeast eIF2B in the crystal. Alternatively, it is possible that ISRIB stabilized an active conformation of eIF2B that entailed changes in the disposition of the invisible C-terminal HEAT repeats of the catalytic ε subunit or a conformation which, though relatively enriched, remained too under-populated in the ensemble of cryo-EM particles for detection or was selectively under-represented in the images available for analysis.

It is intriguing however to ignore the potential for species divergence in structure and consider the crystallized yeast eIF2B as the *apo* structure and the cryo-EM-derived model of human eIF2B as the ISRIB-bound, active conformation. This perspective suggests a major difference in the disposition of the largely invisible N-terminal extension of the δ subunit between the yeast (*apo* structure) and the ISRIB-ligated human structure. The region of the human δ subunit implicated contains two residues (R170 and V177) that are distant from the ISRIB binding pocket, but nonetheless important to ISRIB action (hamster δ R171 and V178, reported in Sekine et al., 2015). It is tempting therefore to speculate that an ISRIB-induced change in the disposition of this segment of the δ subunit contributes to the ISRIB-induced increase in eIF2B activity. This might come about if the ISRIB induced conformational change in the disposition of the N-terminal extension of the δ subunit were to propagate through the regulatory core to the cavity formed by the convergence of the tips of the α, β and δ subunits enfeebling binding of the N-terminal portion of eIF2α (Vazquez de Aldana and Hinnebusch, 1994; Pavitt et al., 1997; Kashiwagi et al., 2016; Bogorad et al., 2017). The region surrounding the cavity is rather flexible and the low resolution of the cryo-EM density in that region might have masked important allosteric changes (Figure 2-supplement 2A).

ISRIB antagonizes the eIF2(αP)-dependent ISR, but *in vitro* ISRIB enhances eIF2B GEF activity even in the absence of phosphorylated eIF2α (Sekine et al., 2015; Sidrauski et al., 2015b). Therefore, the consequences of any ISRIB-induced conformational change in eIF2B must not be limited to weaker binding of the phosphorylated form of eIF2. A recent study suggests that eIF2B exerts its GEF activity by stabilizing the transitional, nucleotide free (*apo*) state of eIF2 and that the difference in free energy between the interaction of the *apo* and nucleotide-bound eIF2 with eIF2B drives catalysis. Phosphorylation inhibits GEF activity by enhancing the interaction of the N-terminal portion of the α subunit of nucleotide bound eIF2 with the regulatory core of eIF2B, thus diminishing the aforementioned free energy gradient (Bogorad et al., 2017). It is tempting to speculate that by enfeebling the interaction of the nucleotide bound eIF2 with eIF2B, ISRIB binding might enhance the free energy difference to facilitate GEF activity.

The consequences of ISRIB binding may not be limited to its effect on GEF activity, as eIF2B also possess a guanine nucleotide dissociation inhibitor (GDI) – displacement activity, whereby eIF2B promotes the dissociation of a stabilizing complex between eIF5 and the GDP-bound eIF2 (Jennings et al., 2013). Enhancement of eIF2B’s GDI-displacing activity could contribute to ISRIB’s activity as an ISR inhibitor.

The regulatory and catalytic subunits of eIF2B evolved from ligand regulated protein ancestors. ADP-glucose pyrophosphorylase has a nucleotide bind site that is conserved in its descendant catalytic γ and ε subunits, whereas ribose-1,5-bisphosphate isomerase has a phospho-sugar binding pocket conserved in its descendant α, β and δ regulatory subunits (reviewed in Kuhle et al., 2015; Kashiwagi et al., 2017). The ISRIB binding pocket discovered here represents a third conserved feature of eIF2B. It is intriguing to consider that endogenous ligand(s) might exist that engage the ISRIB binding site to regulate eIF2B in yet to be determined physiological states.

## Materials and Methods

### Cell culturing and reagents

HeLa-derived cell line was maintained in DMEM Joklik modification (M0518, Sigma) supplemented with 0.2% NaHCO_3_ (S8761, Sigma), Newborn Calf Serum (N4637, Sigma), 1× Penicillin/Streptomycin (P0781, Sigma), 1× non-essential amino acid solution (M7145, Sigma), and 55 μM β-mercaptothanol. Suspension culture was grown in either Erlenmeyer flasks in a Minitron shaker (Infors HT) at a shaking speed 70 rpm, 37°C, 8% CO_2_, or in a Cellbag (BC10, GE Healthcare) using WAVE Bioreactor 20/50 EHT system (GE Healthcare) at 37°C, 8% CO_2_, air flow 0.3 sp, angle 5°-7.5°, rotation speed 15-18.5 rpm.

CHO-K1-derived cell lines were maintained either in Nutrient Mixture F12 (N4888, Sigma), 10% Fetal Calf serum (FetalClone II, Thermo), 2 mM L-glutamine, and 1× Penicillin/Streptomycin at 37°C with 5% CO_2_ for adherent cells or in MEM Alpha (M8042, Sigma) with 10% Fetal Calf serum, 2 mM L-glutamine, 1x Penicillin/Streptomycin, 1× non-essential amino acid solution, and 55 μM β-mercaptothanol in Erlenmeyer flasks in Minitron shaker (Infors HT) at a shaking speed 70 rpm, 37°C, 8% CO_2_ for suspension cells.

### Sucrose Gradient

75×10^6^ cells (in 75 mL) of HeLa-2C2 (3xFLAG-EIF2B2 in/in) cell suspension were treated for 20 minutes with either 150 μL DMSO or 200 nM ISRIB, prior to harvesting and lysing in 3x cell pellet volume of the lysis buffer [50 mM Tris (pH 7.4), 400 mM KCl, 4mM Mg(OAc)_2_, 0.5% Triton, 5% Sucrose, 1 mM DTT, 2 mM PMSF, 10 μg/ml Aprotinin, 4 μg/μl Pepstatin, 4 μM Leupeptin]. Lysates were cleared at 21,130 g on a chilled centrifuge, 0.5 mL of the supernatant was applied on 5mL of 5-20% Sucrose gradient, that was prepared on SG15 Hoeffer Gradient maker in the cell lysis buffer with respective amounts of sucrose and equilibrated for 24 hours on ice. The gradient was run on a SW50.1 rotor at 40,000 rpm for 14 hours and 20 minutes and was equally fractionated into 13 fractions of 420 μL. Each fraction was then diluted two-fold with lysis buffer without sucrose, proteins were precipitated with 20% TCA for 16 hours at 4°C and pelleted at 21,130 g for 20 minutes in a chilled centrifuge. Protein pellets were washed twice with ice-cold acetone, air-dried, resuspended in 60 μL of alkaline SDS loading buffer and 5 μL of resuspension was run on a 12.5% SDS-PAGE gel. The protein gel was then transferred onto PVDF membrane and immunoblotted, using primary monoclonal mouse anti-FLAG M2 antibody (F1804, Sigma) and secondary polyclonal goat anti-mouse-IR800, followed by scanning on Odyssey imager (LI-COR Biosciences) and image analysis on ImageJ software.

### Protein purification

Human eIF2B was purified from 50 L of HeLa-2C2 cells (3xFLAG-EIF2B2 in/in) grown in suspension at a maximum density of 10^6^ cells/mL. Cell pellets (150 grams total) were harvested at room temperature and washed twice with room temperature PBS. Cells were then lysed in 2 pellet volumes of lysis buffer [50 mM Tris (pH 7.4), 150 mM NaCl, 0.5% Triton, 10% Glycerol, 5 mM MgCl_2_, 1 mM DTT, 2 mM PMSF, 10 μg/ml Aprotinin, 4 μg/μL, Pepstatin, 4 μM Leupeptin]. For structural analysis lysis buffer was supplemented with 200 nM ISRIB. Lysates were cleared at 20,000 rpm (JA 25.50) at 4°C and supernatants were incubated with 5 mL of anti-FLAG M2 affinity gel (A2220, Sigma) for one hour at 4°C. The resin with bound eIF2B was washed three times for 5 minutes with 10 mL of ice-cold lysis buffer containing 500 mM NaCl, then washed three times for 5 minutes with 10 mL of washing buffer [50 mM Tris (pH 7.4), 150 mM NaCl, 0.1% CHAPS, 5m M MgCl2, 1 mM DTT]. The bound protein was eluted twice sequentially in 10 mL of the washing buffer supplemented with 150 mg/mL 3X FLAG peptide (F4799, Sigma), concentrated to 2 mg/mL using 100K MWCO PES concentrator (88503, Pierce) and 100 μL of concentrated protein complex was run on a SEC-3 300 Å HPLC column (P.N. 5190-2513, Agilent) at a flow rate of 0.35 mL/min in gel filtration buffer [50 mM Tris (pH 7.4), 150 mM NaCl, 5 mM MgCl_2_, 1 mM DTT]. The centre of the 280 nm absorbance peak eluting around 6 minutes was collected in 42 μL and then applied onto cryo-grids for subsequent cryo-electron microscopy imaging. For the fluorescence polarization experiments the concentrated FLAG-M2 eluate was used.

Hamster eIF2B from CHO-S7 [*Eif2b3*-3xFLAG in/in; *Eif2b4*^wt^], CHO-S9 [*Eif2b3*-3xFLAG in/in; *Eif2b4*^L180F^ in/Δ] or CHO-B3 [*Eif2b3*-3xFLAG in/in; *Eif2b2*^H188K^ in/ Δ] was purified in a way similar to human eIF2B (Sekine et al., 2015), with following exceptions.

Hamster eIF2B^wt^ cells were harvested from 3.5 L of CHO-S7 suspension culture grown at maximum density of 10^6^ cells/mL, washed with room temperature PBS and lysed in two pellet volumes of lysis buffer. Cleared lysate supernatant was incubated with 300 μL of anti-FLAG M2 affinity gel. The resin was washed three times for 5 minutes with 1 mL of lysis buffer containing 500 mM NaCl, then washed three times for 5 minutes with 1 mL of washing buffer [50 mM Tris (pH7.4), 150 mM NaCl, 2 mM MgCl_2_, 0.01% Triton, 1 mM DTT]. Protein was eluted in two resin volumes of washing buffer supplemented with 125 mg/mL 3X FLAG peptide. Hamster eIF2Bδ^L180F^ along with eIF2Bβ^H188K^ mutant cells (with total cell count of 50×10^7^ each) were harvested and washed with room temperature PBS, then lysed in three pellet volumes of lysis buffer. Cleared lysate supernatant was incubated with 60 μL of anti-FLAG M2 affinity gel following binding, washing and eluting procedures as described above.

### Fluorescence polarization assay

20 μL reactions were mixed in 384-well round bottom polystyrene plates (3677, Corning), equilibrated in the assay buffer [50 mM Tris (pH7.4), 150 mM NaCl, 2 mM MgCl_2_, 0.01% Triton, 1 mM DTT] for 10 minutes at room temperature and either read directly or read in 25 minutes after addition of unlabelled competitor on a CLARIOstar microplate reader (BMG Labtech) with filter settings of 482 nm (excitation) and 530 nm (emission). The FAM-labelled AAA2-101 compound was used in the range of 1-100 nM with corresponding amounts of purified eIF2B of 30-250 nM. Each measurement point was either an average of six readings every 30 seconds (for eIF2B dose-response) or five readings every 60 seconds (for unlabelled competitor assay).

### Generation of the genome edited cells by CRISPR/Cas9

Cell lines HeLa-2C2 cells [3xFLAG-*EIF2B2* in/in], CHO-S7 [*Eif2b3*-3XFLAG in/in; *Eif2b4*^wt^] and CHO-S9 [*Eif2b3*-3xFLAG in/in; *Eif2b4*^L180F^ in/Δ] have been previously described (Sekine et al., 2015).

Site-directed random mutagenesis of hamster *Eif2b2* was carried out using CRISPR/Cas9 homologous repair using an equimolar mixture of single-stranded oligo deoxynucleotides (ssODN) as repair templates containing combination of all possible codons (n=64) for each targeted site proving the diversity of substitutions. Each set of ssODNs was transfected along with a sgRNA guide, inserted into a vector containing mCherry marked Cas9, in the following pairs: UK2105 with ssODN-1922 or ssODN-1923, UK2106 with ssODN-1924 (see plasmid and primer tables for further description of guide vectors (UK2105, UK2106) (Table S4) and repair templates (Table S5).

For transfection 20×10^3^ CHO-S7 cells (Sekine et al., 2015) were plated 36 hours prior being transfected with equal amounts of 1 μg guide and 1 μg ssODN using standard Lipofectamine protocol, expanded and sorted 48 hours later. Preceding phenotypical sorts a technical “mCherry-positive” sort for successfully transfected cells was done and 240K, 360K and 330K cells were collected for N162X, H188X and I190X templates respectively. Five days later the recovered cell pools were treated with 0.5 mM histidinol and 200 nM ISRIB. 20 hours later pools were sorted for “GFP-bright” (ISRIB-resistant) and “GFP-dull” (ISRIB-sensitive) phenotypes (Sort I). 37K, 126K and 240K cells for each of respective ISRIB-resistant pools: N162X, H188X and I190X, and 500K for each of ISRIB-sensitive pools were collected. 10 days later “GFP-dull” sort of untreated pools (Sort II) was carried out, to eliminate clones with a constitutively active ISR. 1 mln cells were collected for each of ISRIB-resistant and ISRIB-sensitive pools.

For a new round of sorting to reselect congener-sensitive pools from ISRIB-resistant *Eif2b*^H188X^ pool, 1 mln cells were plated 36 hours before being treated with 0.5 mM histidinol and 2.5 μM of ISRIB or 2.5 μM ISRIB congener (AAA1-075B or AAA1-084).

20 hours later cells were sorted for “GFP-dull” (congener-sensitive) phenotype, and 1 mln cells were collected for each respective pool (ISRIB-sensitive, 075B-sensitive and 084-sensitive). Note that newly collected ISRIB-resistant pool didn’t undergo any treatments prior being sorted serving as a control population.

Cell pools were then expanded, phenotypically characterized and their genomic DNA obtained for subsequent next-generation sequencing analysis.

Individual clones were genotyped following Sanger sequencing of PCR products amplified from genomic DNA.

All cells were washed with PBS once and collected in PBS containing 4 mM EDTA and 0.5% BSA and sorted on INFLUX cell sorter.

For cell line details see supplementary materials (Table S6).

### NGS sequencing of *Eif2b2* containing targeted mutations

Genomic DNA was prepared from 10^7^ cells using Blood and Cell culture Midi Kit (13343, Qiagen). Then two rounds of PCR were used to isolate amplicons for NGS sequencing. The first round used 18 cycles with primers 1975 and 1976 to amplify a 244 bp region of *Eif2b2* from 31 μg of DNA (corresponding to genomic DNA from 19,300 cells) divided into in 5 PCR reactions of 50 μl using Q5 polymerase (M0493, NEB) according to the manufacturer’s protocol. The PCR reactions were then pooled together and mixed with 250 μL of solubilization buffer from Gene-Jet gel purification kit (K0691, Thermo Fisher). Thereafter mixes were purified following the standard Gene-Jet gel purification protocol (without adding the isopropanol recommended for smaller products). The second round used 11-15 cycles of PCR with a universal Illumina P5 primer (1977) and individually barcoded P7 primers for each sample (1762-1768). The resulting products were purified as above and quantified using Agilent DNA chip 1000 and qPCR with Illumina P5 and P7 primers and subject to Illumina SE150bp NGS sequencing (Figure 3 and tables S1) or SE50 NGS sequencing (Figure 5 and tables S2). Primers details could be found in supplementary materials (Table S5).

The sequencing reads were converted to bam files aligned to the ceEIF2b2 locus and then corresponding sam files were aligned to 31 bp templates surrounding each mutation: N162 (NNNGAXGTGATCATGACCATTGGCTATTCT), H188 (CGAAAGAGGAAGTTCNNNGTCATTGTTGCC) and I190 (CGAAAGAGGAXGTTXCATGTCNNNGTTGCCS). Reads that differed by less than three residues outside the degenerate sequence were counted for codons and amino acids at each position using Python scripts.

### CHOP::GFP reporter assay in CHO-K1 cells

40×10^3^ CHO-K1 cells were plated in 6 well plates. Two days later the culture medium was replaced with 2 mL of fresh medium and compounds added. Immediately before analysis, the cells were washed with PBS and collected in PBS containing 4 mM EDTA. Single cell fluorescent signals (10,000/sample) measured by FACS Calibur (Beckton Dickinson). FlowJo software was used to analyze the data.

### Electron microscopy

3 μL aliquots of the protein complex was applied on glow-discharged holey carbon grids (Quantifoil R2/2). Grids were blotted and flash-frozen in liquid ethane using a Vitrobot automat (Thermo Fisher). Data acquisition was performed under low-dose conditions on a Titan KRIOS microscope (Thermo Fisher) operated at 300 kV. The dataset was recorded on a back-thinned Falcon II detector (Thermo Fisher) at a calibrated magnification of × 80 000 (resulting in a pixel size of 1.75 Å on the object scale) with a defocus range of 2-3.5 μm. An in-house built system was used to intercept the videos from the detector at a speed of 25 frames for the 1.5 seconds exposures. Data were acquired automatically using the EPU software (Thermo Fisher) over one 24 hours session. Summarized information could be viewed in supplementary materials (Table S1).

### Image processing

After visual inspection of the micrographs, 765 images were selected for further processing. The movie frames were aligned with MotionCorr (Li et al., 2013) for whole-image motion correction. Contrast transfer function parameters for the micrographs were estimated using Gctf (Zhang, 2016). 237,486 particles were selected semi-automatically using the e2boxer routine from EMAN2 (Tang et al., 2007). All 2D and 3D classifications and refinements were performed using RELION (Scheres, 2012a, b).

As the *S. pombe* structure was not available at the time, first reference-free 2D classification on a sample of images, followed by a 3D classification step was done using as a reference the crystal structure of the tetrameric eIF2B (βδ)_2_ complex from *Chaetomium thermophilum* (PDB: 5DBO, Kuhle et al., 2015) low-pass filtered to 60 Å, generated an initial model that confirmed the presence of all five subunits in our sample and also the existence of a C2 symmetry.

The whole particle dataset was submitted to 3D classification using the newly generated initial model as a reference and sorted into 8 classes. One selected class, representing 41,750 particles (~ 17 % of the dataset), was used for 3D refinement. To further increase the resolution statistical movie processing was performed as described (Bai et al., 2013). The reported overall resolution of 4.1 Å was calculated using the gold-standard Fourier shell correlation (FSC) 0.143 criterion (Scheres and Chen, 2012) and was corrected for the effects of a soft mask on the FSC curve using high-resolution noise substitution (Chen et al., 2013). The final density map was corrected for the modulation transfer function of the detector and sharpened by applying a negative B factor that was estimated using automated procedures (Rosenthal and Henderson, 2003).

### Model building and refinement

*S. pombe* eIF2B structure (PDB accession code: 5B04, Kashiwagi et al., 2016) was used as a starting model. A poly-alanine (glycines were immediately added to account for flexibility) model was generated and subunits were individually fitted to the density map in Chimera (Pettersen et al., 2004). This model was then finely fitted using real space refinement and loops found to be divergent between *S. pombe* and human were rebuilt in Coot (Emsley et al., 2010). All side chains for which density was clearly resolved (up to 37% of non-Ala non-Gly residues in α and β subunits and as low as 8% in ε subunits) were positioned. The ISRIB molecule was then manually fitted in the density located at the regulatory core of eIF2B.

The model was then refined using phenix.real_space_refine (Adams et al., 2010) to optimize both protein and ligand geometry and limit clashes. Finally, REFMAC in CCP-EM (Burnley et al., 2017) was used to further refine the model and automatically generate the map vs. model FSC curves and validate overfitting (as described by Brown et al., 2015). Briefly, the procedure involved initial random displacement of the atoms within the final model and refinement against one of the two half maps to generate the FSC_work_ curve. A cross-validated FSC_free_ curve was then calculated between this refined model and the other half map. The similarity between FSC_work_ and FSCf_ree_ curves is indicative of the absence of overfitting. Summarized information could be viewed in supplementary materials (Table S1).

### Congeners chemistry

General. LC-MS analyses were performed using a Shimadzu UFLCXR system coupled to an Applied Biosystems API2000 mass spectrometer. The HPLC column used was Phenomenex Gemini-NX, 3μm-110 Å, C18, 50 × 2 mm with a flow rate of 0.5 mL/min and UV monitoring at 220 and 254 nm. Gradient elution: pre-equilibration for 1 minute at 5% eluant B in eluant A; then 5 to 98% B over 2 minutes, 98% B for 2 minutes, 98 to 10% B over 0.5 min, then 10% B for one minute. Eluant A: 0.1% HCOOH in H2O; eluant B: 0.1% HCOOH in MeCN. 1H-NMR spectra were recorded using a Bruker-AV 400 instrument operating at 400.13 MHz and 13C-NMR spectra were recorded at 101.62 MHz. Chemical shifts (δ) are in parts per million (ppm) with reference to solvent chemical shift. High-resolution time-of-flight electrospray (TOF-ES+) mass spectra (HR-MS) were recorded using a Bruker micrOTOF spectrometer.

N,N'-((1r,4r)-Cyclohexane-1,4-diyl)bis(2-(4-chlorophenoxy)acetamide) (2a; trans-ISRIB). This compound was prepared from trans-1,4-cyclohexanediamine (1a) and 4-chlorophenoxyacetyl chloride as described (Sidrauski et al., 2013). 1H-NMR (DMSO-d6): δ 7.97 (d, J = 8.0 Hz, 2H), 7.33 (d, J = 8.8 Hz, 4H), 6.96 (d, J = 8.6 Hz, 4H), 4.44 (s, 4H), 3.58 (br s, 2H), 1.75 (br d, J = 7.5 Hz, 4H), 1.33 (quint, J = 10.2 Hz, 4H); 13C NMR (DMSO-d6): δ 167, 157, 129, 125, 117, 67, 47, 31; LC-MS: m/z 451.3 [M + H]+; HPLC: tR 2.92 min (> 95%).

N,N'-((1s,4s)-Cyclohexane-1,4-diyl)bis(2-(4-chlorophenoxy)acetamide) (2b; cis-ISRIB). This compound was prepared from cis-1,4-cyclohexanediamine (1b) and 4-chlorophenoxyacetyl chloride as described (Sidrauski et al., 2013). ^1^H-NMR (DMSO-d6): δ 7.87 (d, J = 7.1 Hz, 2H), 7.34 (d, J = 9.5 Hz, 4H), 6.98 (d, J = 8.7 Hz, 4H), 4.49 (s, 4H), 3.73 (br s, 2H), 1.54–1.63 (m, 8H); 13C-NMR (DMSO-d6): δ 167, 157, 130, 125, 117, 67, 45, 28; LC-MS: m/z 451.4 [M + H]+; HPLC: tR 2.96 min (95%).

N,N'-((1r,4r)-Cyclohexane-1,4-diyl)bis(2-methoxyacetamide) (3; AAA1-090). A stirred solution of 1a (0.11 g, 1 mmol) and iPr2NEt (0.37 mL, 2.2 mmol) in CH_2_Cl_2_ (5 mL) was cooled to 0^o^C. Methoxyacetylchloride (0.19 mL, 2.2 mmol) in CH_2_Cl_2_ (1 mL) was added dropwise. The resulting suspension was stirred at 0°C for 30 minutes and at room temperature for 1 hour. EtOAc was added and the solution was extracted successively with saturated aqueous solutions of NH_4_Cl, NaHCO_3_ and brine. The organic phase was dried with Na_2_SO_4_, filtered, and evaporated. The residue was triturated with Et_2_O, collected and dried to afford the title compound as a white powder (0.219 g, 85%) ^1^H-NMR (DMSO-d6): δ 7.55 (d, J = 8.8 Hz, 2H), 3.75 (s, 4H), 3.55 (br s, 2H), 3.28 (s, 6H), 1.70 (d, J = 8.4 Hz, 4H), 1.33 (quint, J = 19.7 Hz, 4H); 13C-NMR (DMSO-d6): δ 168, 72, 59, 47, 31; HR-MS: m/z 259.1660 [M + H]+, C_12_H_23_N2O_4_ requires 259.1652.

N-((1r,4r)-4-Aminocyclohexyl)-2-(4-chlorophenoxy)acetamide (4b). A stirred solution of tert-butyl ((1r,4r)-4-aminocyclohexyl)carbamate (1c, 2.143 g, 10 mmol) and iPr_2_NEt (1.9 mL, 11 mmol) in CH_2_Cl_2_ (50 mL) was cooled to 0^o^C (ice bath). 4-Chlorophenoxyacetyl chloride (1.7 mL, 11 mmol) in CH_2_Cl_2_ (3 mL) was added dropwise. The resulting suspension was stirred at 0^o^C for 30 minutes and at room temperature for 1 hour. EtOAc was added and the solution was extracted successively with saturated aqueous solutions of NH_4_Cl, NaHCO_3_ and brine. The organic phase was dried with Na_2_SO_4_, filtered and evaporated. The residue was triturated with Et_2_O, collected and dried to afford the Boc-protected intermediate 4a as a white powder (3.63 g, 95%).

A suspension of 4a (1.43 g, 3.5 mmol) in dioxane–MeOH (2:1; 6 mL) was treated with 4 M HCl in dioxane (7 mL, 28 mmol) with stirring for 2 hours. The resulting solution was evaporated and the residue was triturated with Et_2_O, collected and dried to afford the HCl salt of the title compound 4b as a white powder (1.071 g, 96%). ^1^H-NMR (DMSO-d6): δ 8.0 (d, J = 6.5 Hz, 1H), 7.33 (d, J = 8.7 Hz, 2H), 6.96 (d, J= 8.7 Hz, 2H), 5.98 (br s, 2H), 4.46 (s, 2H), 3.50-3.60 (m, 1H), 2.91 (br s, 1H), 1.97, 1.78 (dd, J = 10.9 Hz, J = 13.1 Hz, 4H), 1.36 (sext, 4H); 13C-NMR (DMSO-d6): δ 167, 157, 129, 125, 117, 67, 49, 47, 31, 29; HPLC: tR 2.03 min (95%).

6-chloro-N-((1r,4r)-4-(2-(4-chlorophenoxy)acetamido)cyclohexyl)chromane-2-carboxamide (6; AAA1-084). To a stirred solution of (rac)-6-chlorochromane-2-carboxylic acid (5; 0.1 g, 0.5 mmol; prepared as described (Dolle, 2005) and Et_3_N (0.13 mL, 1 mmol) in CH_2_Cl_2_ (5 mL) was added O-(benzotriazol-1-yl)-N,N,N’,N’-tetramethyluronium hexafluorophosphate (HBTU; 0.21 g, 0.6 mmol). The mixture was stirred for 10 min, when 4b (0.16 g, 0.55 mmol) was added. Stirring was continued overnight, the reaction was diluted with H_2_O, and was extracted with EtOAc. The organic layer was washed with aqueous NH_4_Cl and brine, dried over Na_2_SO_4_, filtered and evaporated. The residue was triturated with Et_2_O, collected and dried to afford the title compound as an off-white powder (0.155 g, 65%). ^1^H-NMR (DMSO-d6): δ 7.95 (d, J = 8.2 Hz, 1H), 7.83 (d, J = 7.5 Hz, 1H), 7.35 (d, J = 8.3, 2H), 7.14 (d, J = 7.2, 2H), 6.97 (d, J = 9.3, 2H), 6.89 (d, J = 9.0 Hz, 1H), 4.51, 4.49 (dd, J = 3.9 Hz, J = 3.0 Hz, 1H), 4.46 (s, 2H), 3.59 (br s, 2H), 2.84-2.76 (m, 1H), 2.72-2.65 (m, 1H), 2.17-2.10 (m, 1H), 1.90-1.82 (m, 1H), 1.78 (br s, 4H), 1.40-1.30 (m, 4H); 13C-NMR (DMSO-d6): δ 169, 168, 157, 152, 129, 126, 125, 124, 118, 117, 75, 67, 48, 31, 25, 23; HPLC: tR 3.01 min (95%). HR-MS: m/z 477.1350 [M + H]+, C_24_H_27_Cl_2_N_2_O_4_ requires 477.1342.

6-Chloro-N-((1r,4r)-4-(2-(4-chlorophenoxy)acetamido)cyclohexyl)-3,4-dihydro-2H-benzo[b][1,4]oxazine-2-carboxamide (8a; AAA1-075B). A solution of (rac)-ethyl 6-chloro-3,4-dihydro-2H-1,4-benzoxazine-2-carboxylate (7a; 0.25 g, 1 mmol, prepared as described) (Carr, 1979) in THF–H_2_O (2:1) was treated with LiOH.H_2_O (0.12 g, 3 mmol) and the resulting mixture was stirred for 3 hours. The solution was neutralized with 1 M aq HCl solution and extracted with EtOAc. The organic phase was dried over Na_2_SO_4_, filtered and concentrated to afford 6-chloro-3,4-dihydro-2H-1,4-benzoxazine-2-carboxylic acid 7b (0.19 g, 89%).

To a stirred solution of 7b (0.15 g, 0.7 mmol) and Et_3_N (0.18 mL, 1.4 mmol) in CH_2_Cl_2_ (5 mL) was added HBTU (0.3 g, 0.84 mmol). The mixture was stirred for 10 min, when 4b (0.23 g, 0.77 mmol) was added. Stirring was continued overnight, the reaction was diluted with H_2_O, and was extracted with EtOAc. The organic layer was washed with aqueous NH_4_Cl and brine, dried over Na_2_SO_4_, filtered and evaporated. The residue was dissolved in MeOH (1 mL) and the product was precipitated by adding an excess of Et_2_O. The precipitate was collected, washed with more Et_2_O and dried to afford the title compound as an off-white powder (0.224 g, 67%). ^1^H-NMR (DMSO-d6): δ 8.08 (d, J = 8.4 Hz, 1H), 7.92 (d, J = 8.4 Hz, 1H), 7.34 (d, J = 8.7 Hz, 2H), 6.97 (d, J = 9.0 Hz, 2H), 6.78 (d, J = 8.4 Hz, 1H), 6.63 (d, J = 2.7 Hz, 1H), 6.50, 6.48 (dd, J = 3.3 Hz, J = 2.7 Hz, 1H), 6.31 (s, 1H), 4.47 (s, 2H), 4.44, 4.42 (dd, J = 3.3 Hz, J =3.6 Hz, 1H), 3.57 (br s, 2H), 3.45, 3.42 (tt, J = 6.7 Hz, J = 6.2 Hz, 1H), 3.20-3.15 (m, 1H), 1.78-1.70 (m, 4H), 1.42-1.29 (m, 4H); 13C-NMR (DMSO-d6): δ 168, 166, 157, 141, 136, 129, 125, 118, 116, 114, 73, 68, 48, 42, 31; HPLC: tR 2.90 min (95%). HR-MS: m/z 478.1286 [M + H]+, C23H26Cl2N3O4 requires 478.1295.

4- and 5-((2-(2-(2-(2-(6-chloro-2-(((1r,4r)-4-(2-(4-chlorophenoxy)acetamido)cyclohexyl)carbamoyl)-2,3-dihydro-4H-benzo[b][1,4]oxazin-4-yl)acetamido)ethoxy)ethoxy)ethyl)carbamoyl)-2-(6-hydroxy-3-oxo-3H-xanthen-9-yl)benzoic acid (8f; AAA2-101). To a solution of 8a (0.19 g, 0.4 mmol) in DMF (5 mL) was added K_2_CO_3_ (0.18 g, 1.2 mmol), NaI (0.09 g, 0.6 mmol), and methyl bromoacetate (0.1 mL, 0.48 mmol). The mixture was stirred at 125°C for 16 hours, cooled to room temperature and diluted with H_2_O (10 mL). The solution was extracted with EtOAc and the organic phase was washed with H2O and brine, dried over Na_2_SO_4_, filtered and evaporated. The residue was purified by silica gel column chromatography (CH_2_Cl_2_–THF gradient elution) to afford methyl ester 8b (0.14 g, 65%).

A solution of 8b (0.1 g, 0.18 mmol) in THF–H_2_O (2:1) was treated with LiOH.H_2_O (0.037 g, 0.9 mmol) and the resulting mixture was stirred for 3 hours. The reaction was neutralized with 1 M aq HCl and extracted with EtOAc. The organic phase was dried over Na_2_SO_4_, filtered, and evaporated to afford the carboxylic acid derivative 8b (0.09 g, 98%).

This material (0.09 g, 0.2 mmol) and Et_3_N (0.06 mL, 0.5 mmol) were dissolved in CH_2_Cl_2_ (3 mL). HBTU (0.08 g, 0.22 mmol) was added and the mixture was stirred for 10 minutes, when tert-butyl (2-(2-(2-aminoethoxy)ethoxy)ethyl)carbamate (0.054 g, 0.22 mmol) in CH_2_Cl_2_ (1 mL) was added. The mixture was stirred overnight, diluted with H_2_O and extracted with EtOAc. The organic phase was washed with aqueous NH_4_Cl and brine, dried over Na_2_SO_4_, filtered and evaporated. The residue was purified by silica gel column chromatography (CH_2_Cl_2_–MeOH gradient elution) to afford the Boc-protected intermediate 8d (0.12 g, 80%).

A solution of 8d (0.12 g, 0.15 mmol) in dioxane (0.5 mL) was treated with 4 M HCl in dioxane (1 mL, 4 mmol). After stirring for 2 hours the solution was evaporated. The residue was dissolved in MeOH (0.5 mL) and the product was precipitated by adding excess Et_2_O. The precipitate was collected and dried to afford the HCl salt of the amine derivative 8e (0.10 g, 98%).

This material (14 mg, 0.02 mmol) and 5-(and 6-) carboxyfluorescein succinimidyl ester (9.4 mg, 0.02 mmol; NHS-fluorescein, 5/6-carboxyfluorescein succinimidyl ester mixed isomer (46410, ThermoFisher Scientific), were dissolved in DMF (1 mL) with Et_3_N (4 μL). The mixture was stirred overnight at 31^o^C and was evaporated. The residue was fractionate by semi-preparative HPLC (Phenomenex Gemini 5 μm-110 Å, C18, 150 × 10 mm; 5 mL/min flow rate) with linear gradient elution of H_2_O–MeCN (containing 0.1% HCOOH) from 85:15 to 5:95 over 13 minutes to afford the title compound 8f as an orange powder (0.015 g, 75%). ^1^H NMR (DMSO-d6): δ 8.88, 8.74 (tt, J = 5.8 Hz, J = 11.7 Hz, 1H), 8.46 (s, 1H), 8.30 (br s, 1H), 8.22, 8.15 (dd, J = 8.6 Hz, J = 8.0 Hz 1H), 8.05-8.11 (m, 1H), 7.96 (t, J = 8.1 Hz, 2H), 7.68 (s, 1H), 7.34 (t, J = 9.7 Hz, 2H), 6.95 (d, J = 8.6 Hz, 2H), 6.82 (d, J = 8.6 Hz, 1H), 6.67 (s, 2H), 6.58 (d, J = 8.9 Hz, 2H), 6.54 (s, 2H), 6.49 (d, J = 8.0 Hz, 1H), 4.61 (quint, J = 4.0 Hz, 1H), 4.44 (s, 2H), 4.44, 3.81-3.96 (m, 4H), 3.5-3.57 (m, 8H), 2.07 (s, 2H), 1.67-1.78 (m, 4H), 1.29-1.39 (m, 4H), 1.23 (s, 4H); HPLC: tR 2.81 min (95%); HR-MS: m/z 1024.2920 [M + H]+, C_52_H_52_C_l2_N_5_O_13_ requires 1024.2933.

**Scheme S1.**
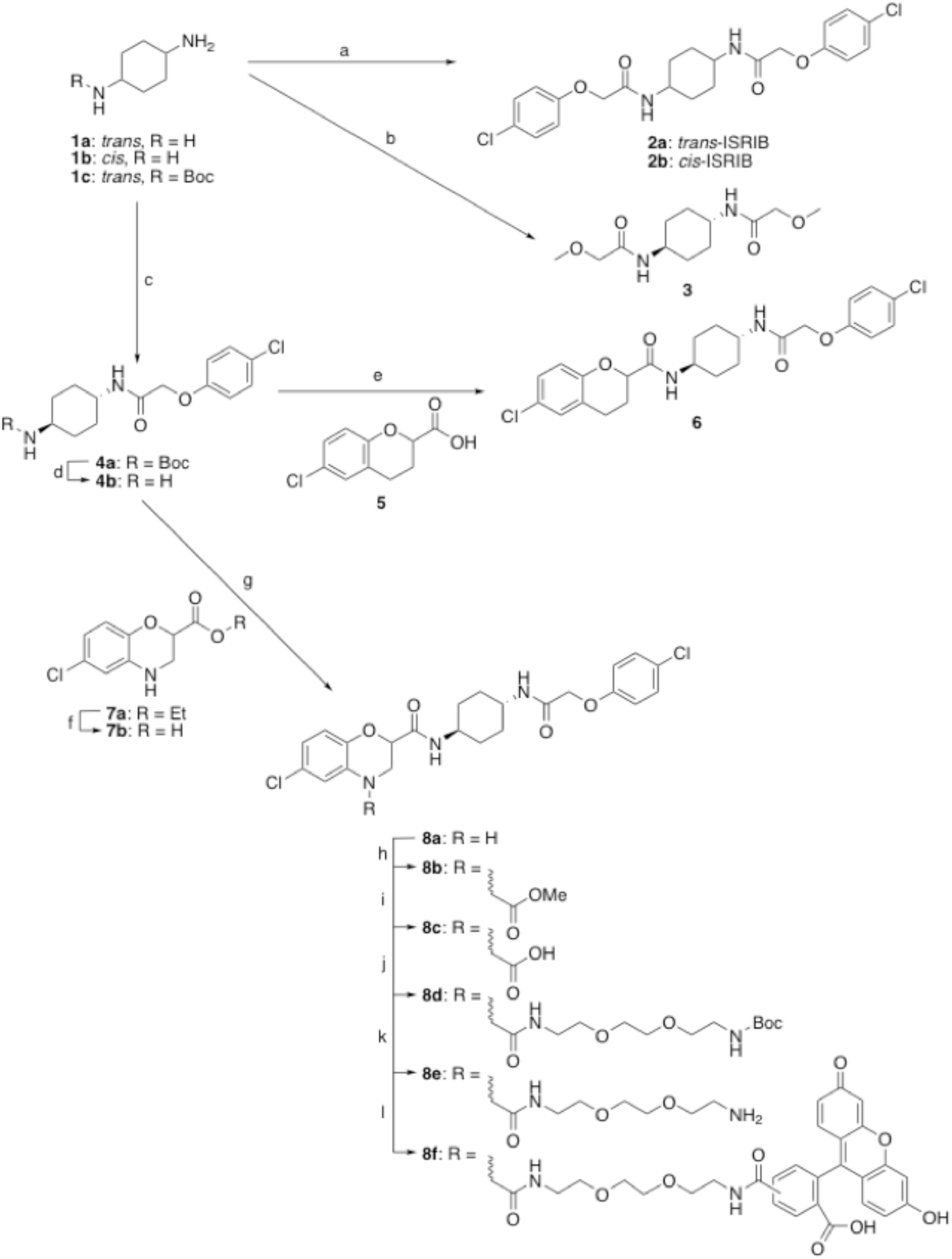
(a) 1a or 1b, 4-chlorophenoxyacetyl chloride, iPr_2_NEt, CH_2_Cl_2_, 0^o^C to rt (70-88%); (b) 1a, methoxyacetyl chloride, iPr_2_NEt, CH_2_Cl_2_, 0°C to rt (85%); (c) 1c, 4-chlorophenoxyacetyl chloride, iPr_2_NEt, CH_2_Cl_2_, 0°C to rt (95%); (d, f, and k) 4 M HCl in dioxane–MeOH (89-98%); (e) 4b and 5, HBTU, Et3N, CH2Cl2 (65%); (g) 4b and 7b, HBTU, Et_3_N, CH_2_Cl_2_ (67%); (h) methyl bromoacetate, K_2_CO_3_, NaI, DMF, 125 oC, 16 hours (65%); (i) LiOH, THF–H_2_O (98%); (j) tert-butyl (2-(2-(2-aminoethoxy)ethoxy)ethyl)carbamate, HBTU, Et_3_N, CH_2_Cl_2_ (80%); (l) 5-(and 6-) carboxyfluorescein.

## Acknowledgements

We thank Dave Barrett and Catherine Ortori (University of Nottingham) for measuring the ISRIB content of eIF2B and Peter Sterk (University of Cambridge) for NGS analysis. We thank S. Chen, C. Savva, S. De Carlo, S. Welsch, F. De Haas, M. Vos and K. Sader for technical support with cryo-EM; G. McMullan for help with movie data acquisition; T. Darling and J. Grimmett for help with computing

Supported by a Wellcome Trust Principal Research Fellowship to D.R. (Wellcome 200848/Z/16/Z) and a Wellcome Trust Strategic Award to the Cambridge Institute for Medical Research (Wellcome 100140). A.A.A. is supported by The Higher Committee for Education Development, Iraq, through a Scholarship (Sponsorship Reference 4241047). AJW is supported by a Specialist Programme from Bloodwise (12048), the UK Medical Research Council (MRC) (MC_U105161083) and a core support grant from the Wellcome Trust and MRC to the Wellcome Trust Medical Research Council Cambridge Stem Cell Institute.

## Author contributions

A.Z. Conceived and led the project; designed and conducted most experiments; analyzed and interpreted the data; prepared the figures and drafted the manuscript.

F.W. Co-designed protein purification for structural analysis; prepared cryo-EM grids; collected and processed cryo-EM data.

A.F. Constructed the atomic model of ISRIB-bound eIF2B; interpreted and analized structural basis of ISRIB binding; prepared figures and edited manuscript.

A.A.A. Designed and synthesized compounds.

A. C-C. Co-conducted and analyzed cell-based assay data on ISRIB derivatives.

H.P.H. Co-designed and supervised somatic cell genetics and deep sequencing analysis; edited the manuscript.

F.A. Analyzed NGS sequencing data and edited manuscript.

L.P. Analyzed NGS sequencing data and edited manuscript.

C.F. Designed and synthesized compounds.

P.F. Designed and synthesized compounds; interpreted the chemogenetic data; edited manuscript.

A.J.W Oversaw the structural analysis of ISRIB binding; edited manuscript.

D.R. Oversaw the project conception and design; interpreted the data; drafted and revised the manuscript.

---

**Figure 1 - Supplement 1.**
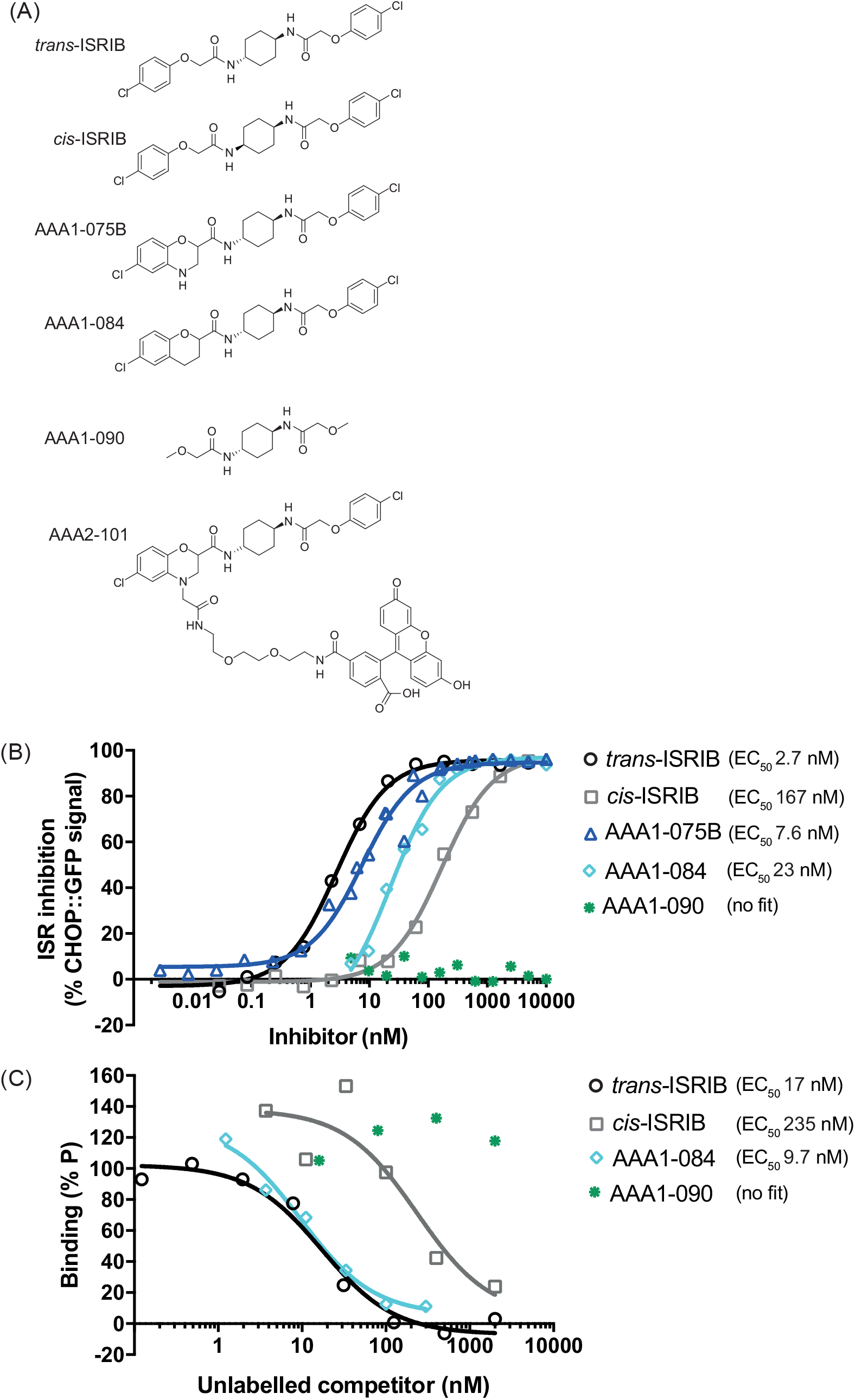
(A) Structures of ISRIB and ISRIB congeners used in this study: active trans-ISRIB, cis-ISRIB, the reference congeners AAA1-075B and AAA1-084, the inactive AAA1-090 and FAM labelled AAA2-101 used in the fluorescence polarization experiments. (B) Plot of the inhibition of the Integrated Stress Response (ISR) activated CHOP::GFP signal, induced upon treatment of CHO-K1 cells with histidinol (HIS) (0.5 mM), in the presence of escalating concentrations of ISRIB (trans-ISRIB) or its congeners. Shown is a representative from three independent experiments conducted for trans and cis-ISRIB and of single experiment conducted for congeners. Concentration of inhibitor is represented on a log_10_ scale. Curve fitting and EC_50_ was generated using “agonist vs. response” function on GraphPad Prism. (C) Plot of the FP signal arising from samples with FAM-labelled AAA2-101 (100 nM) bound to purified human eIF2B (250 nM) in the presence of the indicated concentration of unlabelled ISRIB congeners introduced as competitors. Shown is a representative from three independent experiments. Concentration of unlabelled competitor is represented on a log_10_ scale. Curve fitting and EC_50_ was generated using “agonist vs. response” function on GraphPad Prism.

**Figure 2 - Supplement 1.**
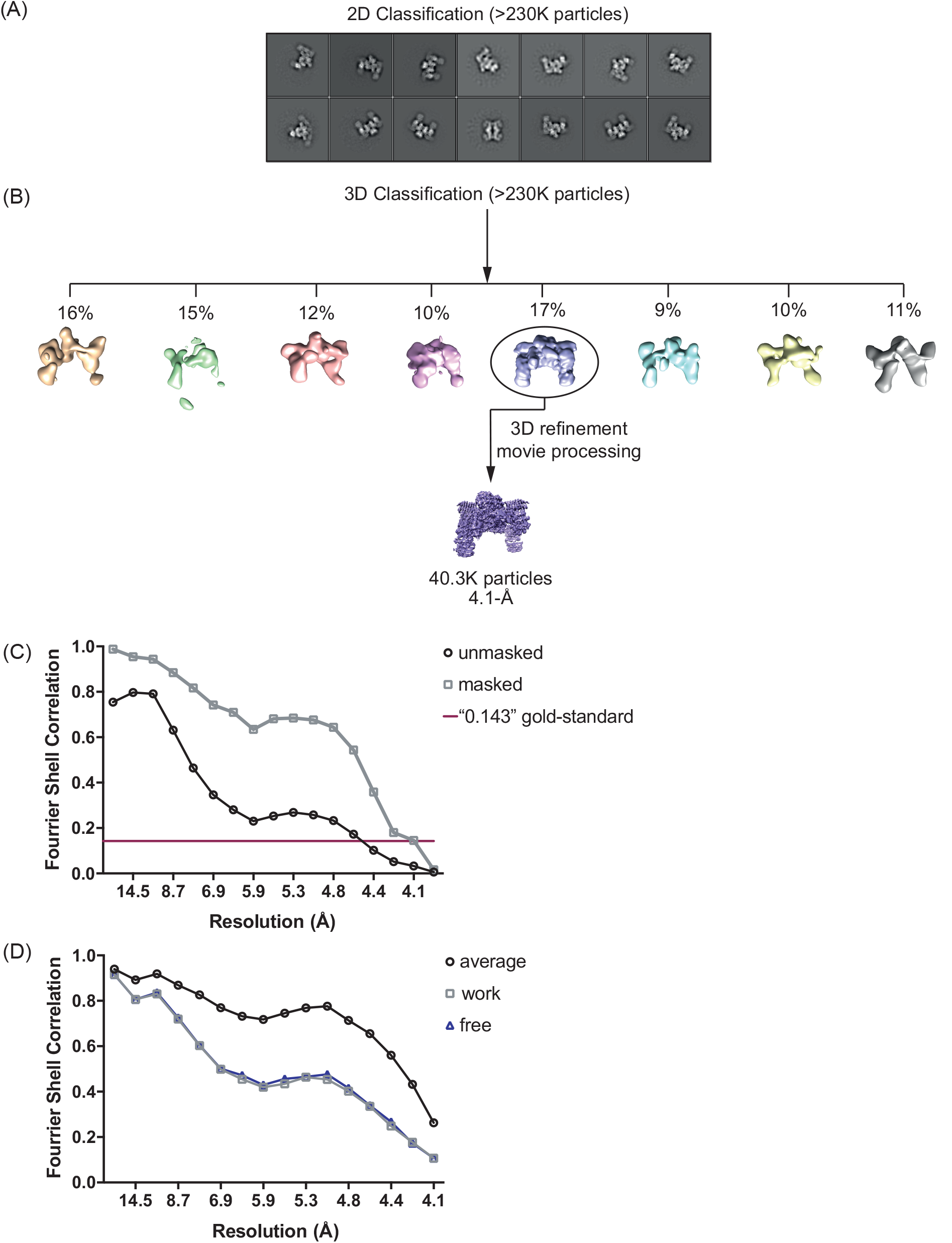
Cryo-electron microscopy data processing. (A) Representative sample of the reference-free 2D class averages of eIF2B particles used for 3D classification. Analysis was performed using RELION and EMAN2 software. (B) 3D classification was conducted on all 2D aligned particles (237,213), 17% (40,326 particles) were used to generate and refine the final 3D map. Analysis was performed using RELION. (C) Gold-standard Fourier shell correlation (FSC) curves. Unmasked map FSC curve is represented in black. Masked-corrected map FSC curve is represented in grey and crosses the “0.143” gold standard purple line at a resolution of 4.1 Å indicating the resolution of the masked map. (D) Refinement and cross-validation of eIF2B model. In black is represented the final map vs. model ‘FSC average’ curve. For cross-validation, the model was ‘shaken’ (see Methods section) and refined against one of the two half-maps to generate the ‘FSC work’ curve (grey line). The ‘FSC free’ curve was calculated between this same refined model and the other half-map (blue line). Superimposition of the two FSC curves shows no sign of over-fitting.

**Figure 2 - Supplement 2.**
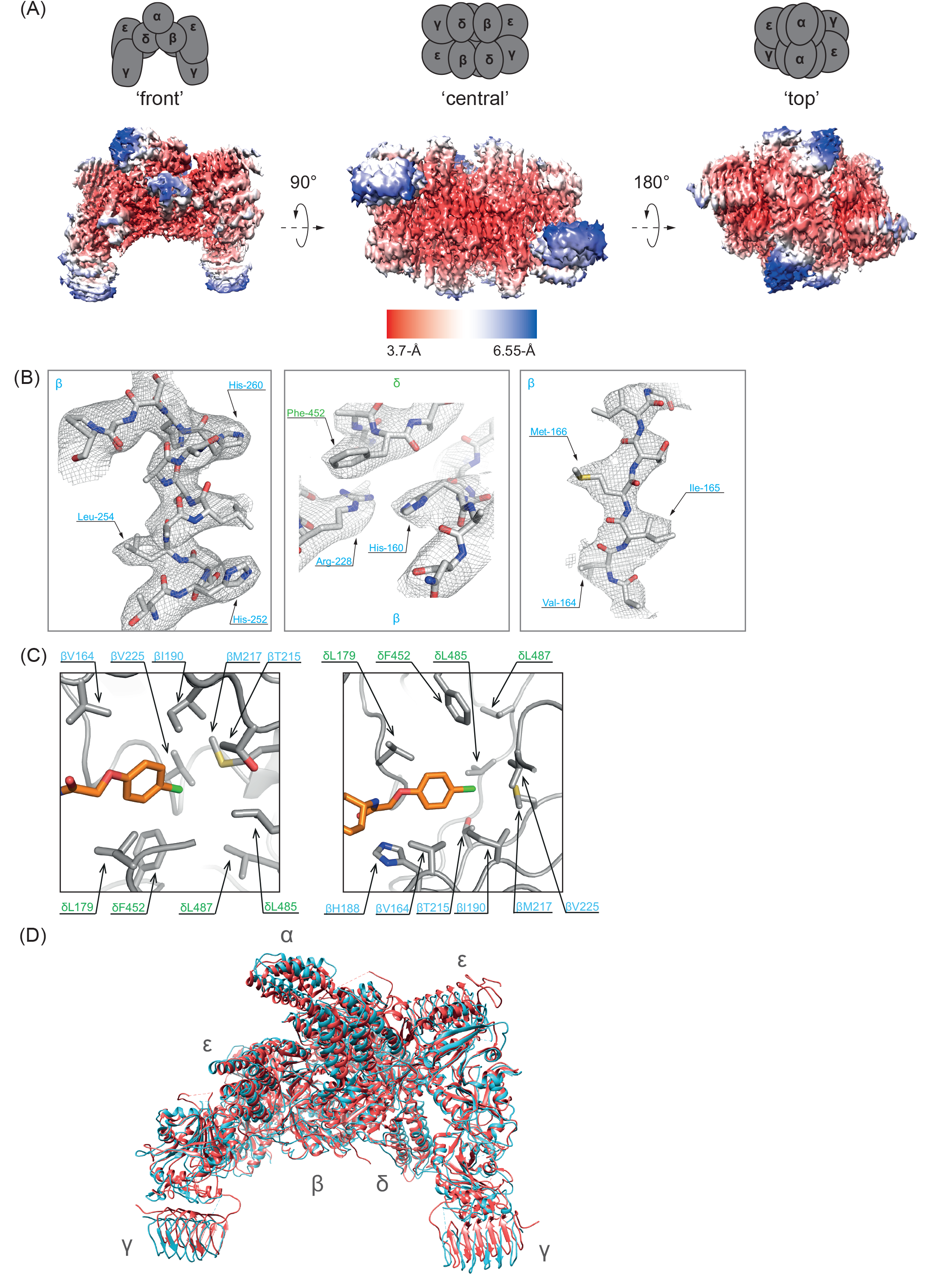
Structure of the ISRIB-bound human eIF2B. (A) Density map colored according to local resolution for the ligated eIF2B complex. The highest resolution is at the core of the (βδ)2 dimer interface, where the putative density of ISRIB was detected. The red-blue palette covers resolution range from 3.7A to >6.55A. (B) Representative density depicted as mesh around the built eIF2B model including well-resolved side chains from the core of (βδ)2 dimer interface. Individual residues of β (blue) and δ (green) subunits are labelled for reference. (C) Close-up of the hydrophobic end of the pocket. Two views of the eIF2B complex core, showing the symmetrically opposite sides of the pocket. The left panel view is from the solvent area looking at the core of the protein. The right panel is from the core looking at the solvent area. (D) Superimposition of ligated human (cyan) and non-ligated yeast eIF2B (PDB: 5B04, red) from the “front view”.

**Figure 4 – Supplement 1.**
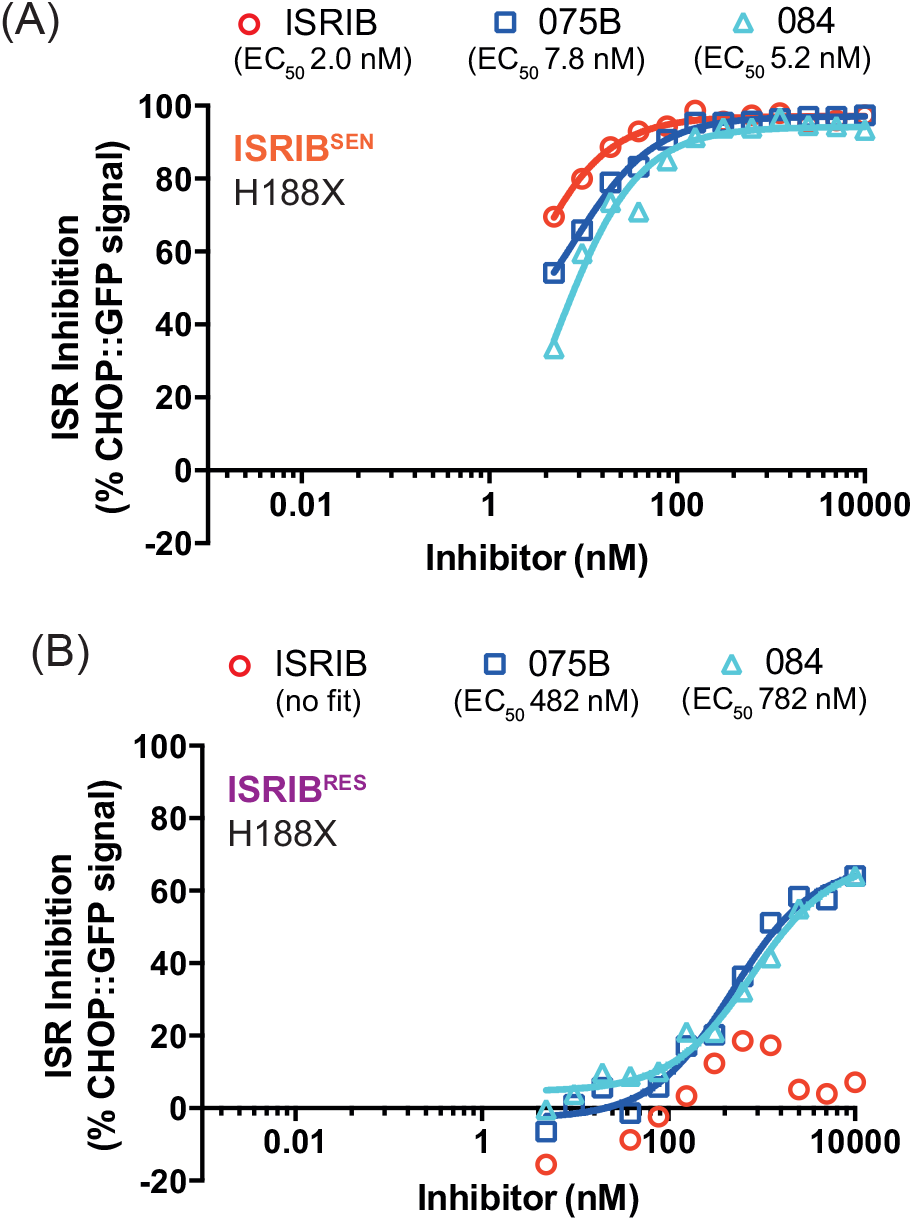
Reproducibility of the effect of mutations in Eif2b2 codon 188 (β^H188X^) on selectivity for ISRIB congeners. Graphs showing inhibition of the ISR-activated CHOP::GFP signal (induced upon treatment with HIS, 0.5 mM) by ISRIB and two related derivatives, AAA1-075B (75B) and AAA1-084 (84), in ISRIB^SEN^ (A) and ISRIB^RES^ (B) mutant pools of *EIf2b2^H188X^*, revealed by flow cytometry. Shown is a representative from three independent experiments for each of the compounds (replicated in Fig. 4 A, B). Concentration of inhibitor is represented on a log_10_ scale. Curve fitting and EC_50_ was generated using “agonist vs. response” function on GraphPad Prism.

**Figure 6 – Supplement 1.**
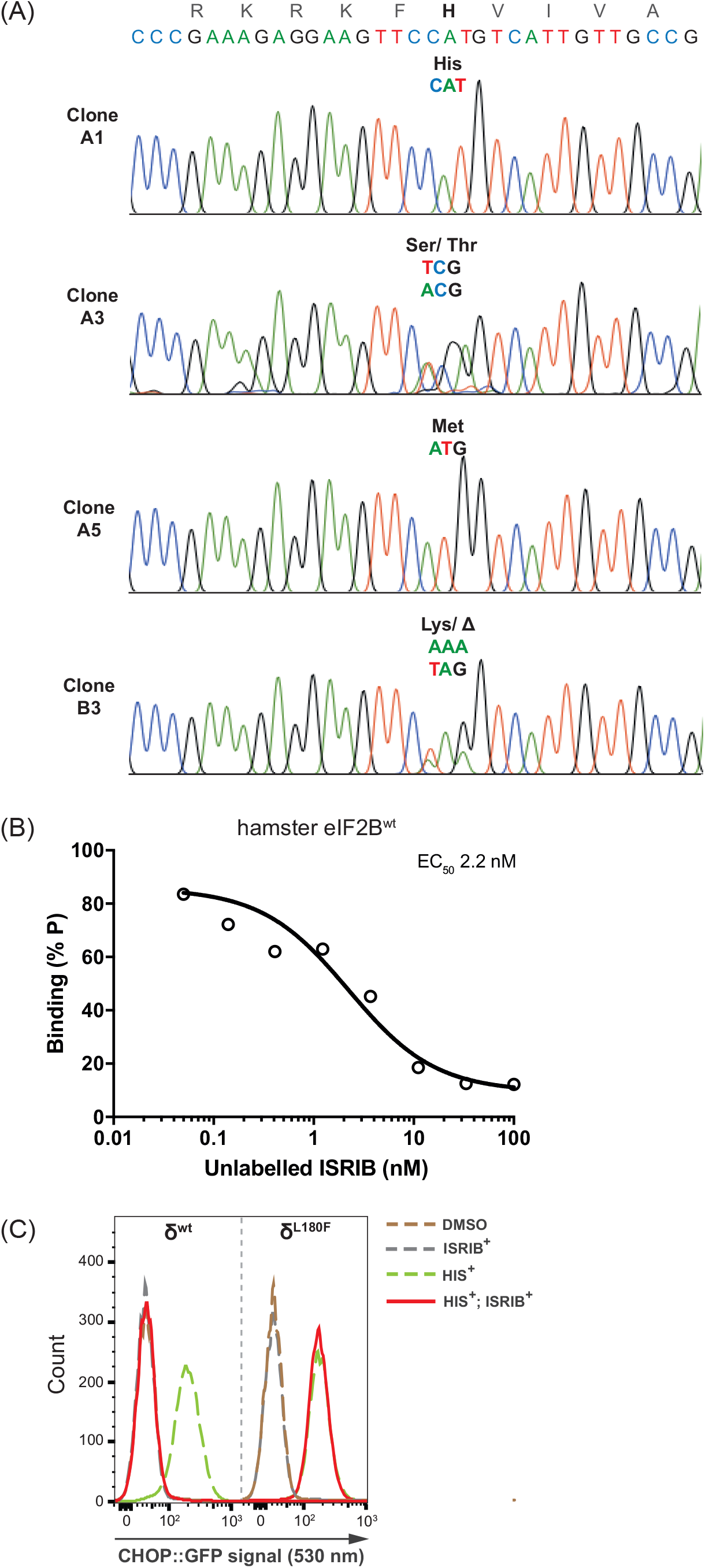
Individual clones from 84^SEN^ pool recapitulate the variation in sensitivity to ISRIB and AAA1-084 observed in the pools. (A) Traces of dye-termination sequencing reactions of PCR amplified *Eif2b2* exon 3 from indicated cell clones: A1 (parental, wt), A3 (H188S/T), A5 (H188M), B3 (H188K/Δ). H188 is highlighted. (B) A plot of the FP signal arising from samples with FAM-labelled AAA2-101 (2.5 nM) bound to purified hamster eIF2B (30 nM) in the presence of the indicated concentration of unlabelled ISRIB introduced as a competitor. Shown is a representative of two independent experiments. Concentrations of ISRIB are represented on a log_10_ scale. The fitting curve and EC_50_ was generated using “agonist vs. response” function on GraphPad Prism. (C) Histograms of CHOP::GFP activity in ISRIB^SEN^ (wt) and ISRIB^RES^ *EIf2b4^L180F^* (clone S9, from Sekine et al., 2015) revealed by flow cytometry. Each pool was treated as indicated with HIS (0.5 mM), ISRIB (200 nM) or both. Note the difference between wt and δ^L180F^ in their ability to inhibit the ISR-activated CHOP::GFP signal when treated with ISRIB (compare continuous red trace in the left and right panels).

**Table S1.** *Mode I refinementandstatistics*

| **Cryo-electron microscopy data collection** |  |
| --- | --- |
| Electron microscope | Titan Krios 300 kV |
| Detector | FEI Falcon II |
| Pixel size (Å) | 1.75 |
| Defocus range (µm) | -2.5 to -3.5 |
| Electron dose (e<sup>-</sup>.Å<sup>-2</sup>.s<sup>-1</sup>) | 30 |
| **Reconstruction** |  |
| Particles used for 3D refinement | 40300 |
| Cell dimensions |  |
| a, b, c (Å) | 240, 240, 240 |
| α, β, γ (°) | 90, 90, 90 |
| **Model refinement** |  |
| Resolution limit | 4.1 |
| Map sharpening factor (Å<sup>2</sup>) | -140.4 |
| FSC<sub>average</sub><sup>1</sup> | 0.6931 |
| RMS bonds deviation (Å) | 0.0069 |
| RMS angles deviation (Å) | 1.3485 |
| **Model validation** |  |
| Molprobtity score | 2.33 (99th percentile) |
| Clashscore | 6.84 (100th percentile) |
| Ramachandran outliers | 0.75% |
| Sidechain outliers | 0.20% |

<sup>1</sup> FSC<sub>average</sub> = $\Sigma(N_{\text{shell}} \text{FSC}_{\text{shell}}) / \Sigma(N_{\text{shell}})$ , where FSC<sub>shell</sub> is the FSC in a given shell, N<sub>shell</sub> is the number of 'structure factors' in the shell.

**Table S2.** *Allele frequency (supplement to figure 3D)*

| Amino Acid | N162X |  |  |  | H188X |  |  |  | I190X |  |  |  |
| --- | --- | --- | --- | --- | --- | --- | --- | --- | --- | --- | --- | --- |
|  | ISRIB-SEN |  | ISRIB-RES |  | ISRIB-SEN |  | ISRIB-RES |  | ISRIB-SEN |  | ISRIB-RES |  |
|  | Reads | % Reads | Reads | % Reads | Reads | % Reads | Reads | % Reads | Reads | % Reads | Reads | % Reads |
| * | 855 | **0.20** | 6575 | **1.74** | 1025 | **0.31** | 6215 | **2.48** | 340 | **0.09** | 3467 | **1.23** |
| A | 3953 | **0.92** | 1768 | **0.47** | 2248 | **0.69** | 4305 | **1.72** | 1106 | **0.31** | 1213 | **0.43** |
| C | 2429 | **0.57** | 1654 | **0.44** | 5992 | **1.83** | 716 | **0.29** | 1416 | **0.39** | 3145 | **1.12** |
| D | 3953 | **0.92** | 1436 | **0.38** | 662 | **0.20** | 5075 | **2.03** | 419 | **0.12** | 5912 | **2.10** |
| E | 2655 | **0.62** | 867 | **0.23** | 227 | **0.07** | 8902 | **3.55** | 426 | **0.12** | 574 | **0.20** |
| F | 2064 | **0.48** | 2798 | **0.74** | 8730 | **2.66** | 682 | **0.27** | 3363 | **0.93** | 10199 | **3.62** |
| G | 419 | **0.10** | 26632 | **7.05** | 965 | **0.29** | 52762 | **21.05** | 949 | **0.26** | 5716 | **2.03** |
| H | 9410 | **2.20** | 10487 | **2.78** | 269253 | **82.06** | 6443 | **2.57** | 208 | **0.06** | 1315 | **0.47** |
| I | 2804 | **0.66** | 14912 | **3.95** | 4980 | **1.52** | 1749 | **0.70** | 326077 | **90.10** | 39307 | **13.96** |
| K | 3520 | **0.82** | 2809 | **0.74** | 499 | **0.15** | 20494 | **8.18** | 139 | **0.04** | 359 | **0.13** |
| L | 10772 | **2.52** | 7055 | **1.87** | 1794 | **0.55** | 46099 | **18.39** | 8405 | **2.32** | 5827 | **2.07** |
| M | 1211 | **0.28** | 724 | **0.19** | 350 | **0.11** | 4795 | **1.91** | 192 | **0.05** | 68523 | **24.34** |
| N | 350813 | **82.08** | 165829 | **43.92** | 3058 | **0.93** | 668 | **0.27** | 398 | **0.11** | 770 | **0.27** |
| P | 2485 | **0.58** | 11242 | **2.98** | 425 | **0.13** | 648 | **0.26** | 331 | **0.09** | 947 | **0.34** |
| Q | 5727 | **1.34** | 1589 | **0.42** | 443 | **0.14** | 15550 | **6.21** | 207 | **0.06** | 2403 | **0.85** |
| R | 7597 | **1.78** | 8707 | **2.31** | 411 | **0.13** | 59827 | **23.87** | 548 | **0.15** | 2188 | **0.78** |
| S | 8007 | **1.87** | 9712 | **2.57** | 2208 | **0.67** | 8331 | **3.32** | 829 | **0.23** | 4443 | **1.58** |
| T | 2143 | **0.50** | 99150 | **26.26** | 1737 | **0.53** | 3153 | **1.26** | 452 | **0.12** | 1932 | **0.69** |
| V | 1721 | **0.40** | 2101 | **0.56** | 13035 | **3.97** | 2950 | **1.18** | 15627 | **4.32** | 1749 | **0.62** |
| W | 428 | **0.10** | 142 | **0.04** | 4038 | **1.23** | 618 | **0.25** | 147 | **0.04** | 79154 | **28.11** |
| X | 253 | **0.06** | 209 | **0.06** | 329 | **0.10** | 256 | **0.10** | 63 | **0.02** | 48 | **0.02** |
| Y | 4159 | **0.97** | 1146 | **0.30** | 5704 | **1.74** | 379 | **0.15** | 262 | **0.07** | 42391 | **15.05** |
| Total Reads | 427378 |  | 377544 |  | 328113 |  | 250617 |  | 361904 |  | 281582 |  |

**Table S3.** *Allele frequency (supplement to figure 5C)*

| Amino Acid | H188X ISRIB-RES |  |  |  |  |  |  |  |
| --- | --- | --- | --- | --- | --- | --- | --- | --- |
|  | ISRIB-RES |  | ISRIB-SEN |  | 075B-SEN |  | 084-SEN |  |
|  | Reads | % Reads | Reads | % Reads | Reads | % Reads | Reads | % Reads |
| **\*** | 8309 | **2.64** | 648 | **0.22** | 4001 | **1.88** | 4428 | **1.63** |
| **A** | 5809 | **1.85** | 4595 | **1.56** | 16814 | **7.89** | 23876 | **8.78** |
| **C** | 1346 | **0.43** | 3465 | **1.17** | 1687 | **0.79** | 2635 | **0.97** |
| **D** | 4608 | **1.47** | 797 | **0.27** | 2338 | **1.10** | 3892 | **1.43** |
| **E** | 8292 | **2.64** | 920 | **0.31** | 1228 | **0.58** | 1311 | **0.48** |
| **F** | 1776 | **0.57** | 12004 | **4.06** | 2450 | **1.15** | 3853 | **1.42** |
| **G** | 52131 | **16.59** | 1558 | **0.53** | 10887 | **5.11** | 12264 | **4.51** |
| **H** | 9311 | **2.96** | 226030 | **76.53** | 34473 | **16.18** | 55443 | **20.40** |
| **I** | 2684 | **0.85** | 4081 | **1.38** | 3612 | **1.70** | 4906 | **1.81** |
| **K** | 27535 | **8.76** | 803 | **0.27** | 47373 | **22.24** | 44470 | **16.36** |
| **L** | 67765 | **21.56** | 6498 | **2.20** | 13552 | **6.36** | 12728 | **4.68** |
| **M** | 6690 | **2.13** | 666 | **0.23** | 11051 | **5.19** | 8943 | **3.29** |
| **N** | 772 | **0.25** | 1915 | **0.65** | 1587 | **0.75** | 1816 | **0.67** |
| **P** | 1193 | **0.38** | 1520 | **0.52** | 1038 | **0.49** | 1298 | **0.48** |
| **Q** | 22530 | **7.17** | 1111 | **0.38** | 4152 | **1.95** | 6029 | **2.22** |
| **R** | 71454 | **22.74** | 1397 | **0.47** | 6097 | **2.86** | 7651 | **2.82** |
| **S** | 11841 | **3.77** | 4824 | **1.63** | 31921 | **14.98** | 49558 | **18.23** |
| **T** | 4857 | **1.55** | 3513 | **1.19** | 11650 | **5.47** | 15931 | **5.86** |
| **V** | 3537 | **1.13** | 11438 | **3.87** | 5317 | **2.50** | 7625 | **2.81** |
| **W** | 1177 | **0.37** | 4767 | **1.61** | 1123 | **0.53** | 2019 | **0.74** |
| **X** | 61 | **0.02** | 61 | **0.02** | 48 | **0.02** | 56 | **0.02** |
| **Y** | 613 | **0.20** | 2757 | **0.93** | 654 | **0.31** | 1105 | **0.41** |
| **Total Reads** | **314291** |  | **295368** |  | **213053** |  | **271837** |  |

**Table S4.** *Plasmid list*

Plasmid list
| ID | Plasmid name | Description | Primers used to generate plasmid | Figure |
| --- | --- | --- | --- | --- |
| UK2105 | EIF2B2_CRISPR7_pSpCas9(BB)-2A-mCherry | mCherry-tagged CRISPR/Cas9 plasmid with guide RNA for targeting hamster *Eif2b2* exon 3 bottom strand | Oligo1925<br>Oligo1926 | 2 |
| UK2106 | EIF2B2_CRISPR8_pSpCas9(BB)-2A-mCherry | mCherry-tagged CRISPR/Cas9 plasmid with guide RNA for targeting hamster *Eif2b2* exon 3 top strand | Oligo1927<br>Oligo1928 | 2 |

**Table S5.**
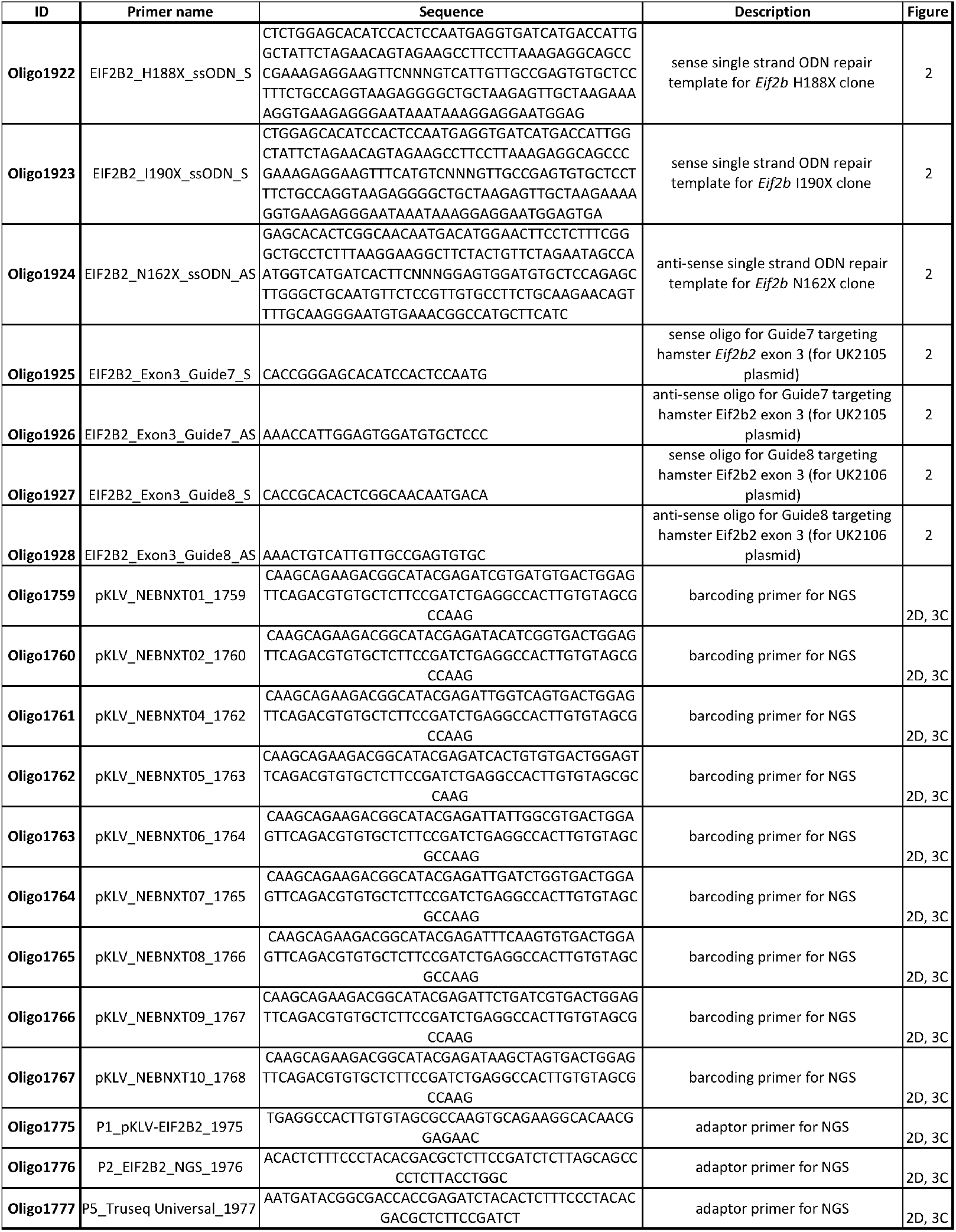
*Oligo list*

**Table S6.** *Cell line list*

| Gene | Exon | Cells | Clone name | Description | Mutagenized region (number shows amino acid position at which mutagenesis occurred) |
| --- | --- | --- | --- | --- | --- |
| *Eif2b2* | 3 | CHO-K1 | **N162X** | mixed population of ISRIB-resistant cells | 162_<u>X</u>EVIMTIGYSRTVEAFLKEAARKRKFHVIV |
| *Eif2b2* | 3 | CHO-K1 | **H188X** | mixed population of ISRIB-resistant cells | 188_<u>X</u>VIVAECAPFCQVRG |
| *Eif2b2* | 3 | CHO-K1 | **I190X** | mixed population of ISRIB-resistant cells | 190_<u>X</u>VAECAPFCQVRG |
| *Eif2b2* | 3 | CHO-K1 | **H188X-075B** | mixed population of 075B-sensitive cells | 188_<u>X</u>VIVAECAPFCQVRG |
| *Eif2b2* | 3 | CHO-K1 | **H188X-084** | mixed population of 084-sensitive cells | 188_<u>X</u>VIVAECAPFCQVRG |
| *Eif2b2* | 3 | CHO-K1 | **A3** | H188S/T, heterozygous, 084-sensitive, mildly ISRIB-resistant | 188_<u>S</u>VIVAECAPFCQVRG;<br>188_<u>T</u>VIVAECAPFCQVRG |
| *Eif2b2* | 3 | CHO-K1 | **A5** | H188M, homozygous, 084-sensitive, strongly ISRIB-resistant | 188_<u>M</u>VIVAECAPFCQVRG |
| *Eif2b2* | 3 | CHO-K1 | **B3** | H188K/Δ, heterozygous, 084-sensitive, strongly ISRIB-resistant | 188_<u>K</u>VIVAECAPFCQVRG |

